# Essential function reflected in the phylodynamics of a multigene family – the *pir* genes of malaria parasites

**DOI:** 10.64898/2026.01.23.697869

**Authors:** Andrew P. Jackson, Deirdre A. Cunningham, Lin Lin, Naomi Mara Claro de Oliveira, Séverine C. Chevalley-Maurel, Giulia Pianta, Franziska Morhing, Abigail K. Renfree, Timothy S. Little, Robert W. Moon, Jean Langhorne, Chris J. Janse, Blandine M.D. Franke-Fayard, Christiaan van Ooij

## Abstract

The genomes of malaria parasites (*Plasmodium spp*.) encode many gene families, which are intimately associated with host interactions and disease in these important pathogens. The largest malaria gene family is the *Plasmodium* interspersed repeat (*pir*) genes, present in rodent, primate and most human malaria parasites, which are suggested to have originated from one highly conserved gene, which we call *pirC1*. The precise function(s) of *pir* is unknown but to determine their potentially multifarious roles we must understand the evolutionary dynamics of *pir* repertoire to discriminate among the many genes. Here we estimate the global phylogeny for *pir* genes in 14 *Plasmodium* species and one *Hepatocystis* species. We reveal that *pirC1* is not the common ancestor but is one of several orthologous genes conserved in multiple species amidst the rapid turnover of species-specific paralogs. We show that the PIRC1 protein is nonetheless essential for blood stage growth of *P. berghei, P. chabaudi* and *P. knowlesi*, as parasites lacking the *pirC1* gene could not be generated or had severely reduced growth rates. As this effect was observed both in vivo and in vitro, the role of *pirC1* is not related to host immune interaction. Rather, *P. berghei* and *P. knowlesi* PIRC1 are secreted from the parasite, pointing to a role in parasite interaction with the host cell or nutrient uptake by blood stages. The phylodynamics of *pir* genes indicate that old orthologs, like *pirC1*, and younger within-species paralogs could have fundamentally different roles, and emphasize the need to distinguish between them in future. This study is the first to provide evidence for the existence of an essential *pir* gene and provides a robust rationale for further experimental approaches to *pir* gene functions.

**SIGNFICANCE:** The genomes of malaria parasites (*Plasmodium*) contain many different gene families, of which the *pir* family is the largest, with more than 1000 members in some species. The PIR proteins are likely important for parasite fitness but their precise functions remain unknown – roles in adherence of infected red blood cells to blood *ves*sels, virulence and immune evasion of have been suggested. How, and why, this highly diverse gene family evolved is a significant question both for understanding malaria physiology and pathogenesis. Here we present a comprehensive *pir* phylogeny, identifying the origins of gene diversity during *Plasmodium* evolution and a select group of highly conserved genes. We show that one conserved *pir* gene (*pirC1*) encodes a protein that is essential for optimal growth of multiple malaria parasites during the asexual blood stage, both in the host and in vitro. This indicates that *pirC1* function relates to interaction with the host cell or nutrient acquisition, and not to immune evasion or sequestration, (although this might still be the function of other *pir* genes). This study provides a robust rationale for the hitherto baffling diversity of *pir* genes, and shows why it is important to distinguish old orthologs from young paralogs in future studies on *pir* gene function.

## INTRODUCTION

Parasites of the genus *Plasmodium* are vector-borne apicomplexan parasites that cause malaria in humans and other vertebrates. In the case of human malaria parasites, the infection invol*ves* three stages: the asymptomatic liver stage, the asexual blood stage (erythrocytic stage), which causes all symptoms of malaria, and the sexual stage, which continues the life cycle when transmitted to mosquitoes. The host-parasite interactions in the blood are mediated by various protein effectors that are often encoded by large parasite-specific gene families (Reid 2015). Examples include families conserved across all *Plasmodium* spp., e.g. the sera genes that encode serine-repeat antigens (Arisue *et al*, 2007) families that are expanded in specific *Plasmodium* species, such as the *fikk* kinase family in *Plasmodium falciparum* (Ward *et al*, 2004; Davies *et al*, 2020), *fam*-a genes in rodent malaria species (Frech & Chen, 2013; Fougère *et al*, 2016), and the var gene family that encodes the EMP1 variant antigens responsible for cytoadherence and immune evasion in the subgenus *Laverania* of *Plasmodium* (i.e. *P(Laverania*)) (Hviid *et al*, 2024). Indeed, multicopy gene families feature are a general phenomenon in most host-parasite interactions, including the *ves* genes in Babesia spp. (Jackson *et al*, 2014), *tpr* genes in *tprTheileria* spp. (Palmateer *et al*, 2023), *vsg* genes in trypanosomes (Silva Pereira *et al*, 2022) and vsp genes in Giardia spp. (Gargantini *et al*, 2016).

The gene family of *Plasmodium* interspersed repeat (*pir*) genes are present in all species of rodent and primate malaria in the subgenera *P(Plasmodium*) and *P(Vinckeia*), but absent from *P(Laverania*) and other subgenera (Janssen *et al*, 2002; del Portillo *et al*, 2001). The number of *pir* genes varies substantially between species and exceeds 1000 in some species (Neafsey *et al*, 2012; Cepeda *et al*, 2024) (Otto *et al*, 2014; Ansari *et al*, 2016). The *pir* loci are commonly distributed in irregular tandem arrays throughout the subtelomeric regions of the chromosomes (Ansari *et al*, 2016; Otto *et al*, 2014; Neafsey *et al*, 2012), although in *P. knowlesi pir*s are found among core chromosomal regions (Pain *et al*, 2008). The subtelomeric location is thought to promote rapid changes in *pir* repertoire through ectopic gene conversion (Cunningham *et al*, 2010) and, indeed, *pir* gene number changes rapidly, varying markedly even between conspecific *Plasmodium* strains (Auburn *et al*, 2016).

The *pir* genes encode transmembrane proteins which, although they have a conserved basic structure (Fig. S1) (Harrison *et al*, 2020), can vary significantly, primarily in length. PIR proteins are expressed throughout the parasite life cycle, particularly during the vertebrate stages from the arrival of the sporozoite in the skin by a mosquito bite, during development in the liver and in both the asexual and gametocyte stages within the erythrocyte (Cunningham *et al*, 2009; Little *et al*, 2021; Otto *et al*, 2014). Different PIRs localise to different cellular locations in the infected host cell, including the parasite cytoplasm and plasma membrane, but some proteins are secreted to the parasitophorous vacuole or exported to the host red blood cell (RBC) cytoplasm or plasma membrane (Yam *et al*, 2016; Rehn *et al*, 2022). Not all PIRs contain obvious export signals and this probably reflects variation in cellular location and function (Cunningham *et al*, 2010).

The function of PIR proteins remains obscure, in part owing to the lack of homology with other proteins (Frech & Chen, 2013). Despite earlier assertions (Janssen *et al*, 2004), recent comparisons of the structures of PIR proteins with RIFIN and STEVOR protein structures revealed that these proteins are not homologous (Harrison *et al*, 2020). The *pir* genes lack the essential characteristics of a variant antigen, such as monoallelic expression, clonality and a lack of heterologous cross-reacting antibodies, indicating that they do not function in classical antigenic variation like PfEMP1 proteins of the var gene family (Fernandez-Becerra *et al*, 2005; Bozdech *et al*, 2008). However, previous studies have suggested some other role in immunity against *Plasmodium* parasites (Cunningham *et al*, 2005, 2009) or virulence (Lin *et al*, 2018). For instance, it has been suggested that PIRs mediate cytoadherence of infected red blood cells in small blood *ves*sels to prevent parasite clearance by the spleen (del Portillo *et al*, 2004), or in red blood cell invasion (Neafsey *et al*, 2012). The sequence diversity and variations in gene expression of *pir* genes, combined with the variety of cellular locations of PIR proteins, may indeed suggest that they perform multiple roles.

The function of *pir* gene has been studied in the human malaria parasites *Plasmodium* vivax and *Plasmodium knowlesi* and the rodent malaria parasites *Plasmodium* berghei, *Plasmodium chabaudi* and *Plasmodium yoelli*, but rarely compared across species. We lack understanding of how *pir* genes relate to each other in the same genome and between genomes of different species, and how relatedness in structure corresponds to function. Classification of *pir* genes (Neafsey *et al*, 2012; Otto *et al*, 2014) has mainly focused on single species (Neafsey *et al*, 2012; Cepeda *et al*, 2024) or closely related parasites (Otto *et al*, 2014). However, there has been no holistic reconstruction of the entire *pir* family, making it difficult to compare studies that analyse *pir* genes in different malaria species. Previous studies have provided evidence for rapid evolution of species-specific (i.e. paralogous) *pir* genes but also for the presence of a highly conserved *pir* gene that occupies a conserved locus in the genomes of multiple *Plasmodium* species (Neafsey *et al*, 2012; Frech & Chen, 2013; Jackson, 2016; Little *et al*, 2021). This particular *pir* gene, which displays both sequence and positional orthology, was named *virD* in *P. vivax* (Neafsey *et al*, 2012), and has been proposed as the ancestor of the entire *pir* gene family (Frech & Chen, 2013; Little *et al*, 2021; Neafsey *et al*, 2012).

In this study, we aim to resolve the phylogeny of *pir* genes in 14 *Plasmodium* genomes and one *Hepatocystis* genome and in*ves*tigate the function of the highly conserved *pir* gene orthologous to *virD*, which we call *pirC1*. Our analysis reveals the origins of *pir* diversity in the common ancestor of primate and rodent *Plasmodium* species and shows how *pir* phylodiversity has fluctuated as these parasites have evolved. We identify multiple clades of orthologous *pir* genes that are widely conserved between species and show in multiple species that one instance (*pirC1*) is essential for growth of blood stages, both in vivo and in vitro. These results are the first genus-wide in*ves*tigation of this gene family and the first direct evidence that a member of a large gene family in *Plasmodium* parasites is essential.

## RESULTS

### Structural differentiation of subfamilies of *pir* genes evolved early

To identify intact, full-length *pir* gene sequences, the genomes of 14 *Plasmodium* and one *Hepatocystis* spp. were searched with BLASTp. The number of *pir* genes obtained ranged from 1179 in P. *ovale* and 794 in *P. cynomolgi* to 12 in *P. inui*, five in *P. fragile* and only four in *Hepatocystis* (Table S1). Although the quality of reference genome sequences varies, it is unlikely that the low number of *pir* genes identified in some genomes are due to incomplete sequencing (SI Results).

The SURFIN proteins have been identified as the closest relati*ves* of the PIR proteins using BLASTp ((Frech & Chen, 2013); SI Results). Therefore, 17 SURFIN sequences from multiple species were included in the multiple alignment. The final alignment of 4232 protein sequences (Data File 1) comprises 837 characters that span the conserved PIR *‘*core*’* and transmembrane domain (del Portillo *et al*, 2001; Janssen *et al*, 2004) (Fig. S1). Variable domains on either side of the transmembrane domain were removed or shortened since these regions differ greatly in length (Fig. S1) and are often only nominally homologous. In 264 PIRs, the conserved core sequence is duplicated or triplicated; for these, only the core sequence duplicate closest to the transmembrane domain was retained (see SI Results for detailed description of the sequence alignment).

A Maximum Likelihood phylogenetic tree was estimated for all sequences using IQ-TREE (Minh *et al*, 2020) on the CIPRES Science Gateway (Miller *et al*, 2010) (Data File 2). A JTT+F+G model of amino acid substitution was found to be optimal and applied with a value for G (alpha) of 3.16. The tree was rooted using outgroup (*surfin*) sequences. The resulting *pir* phylogeny was divided into 23 subfamilies, labeled *pirA* (the most basal-branching clade closest to the root) to *pirW*, with each corresponding to a crown clade of approximately equal age (Fig. 1). These clades are robust, supported by bootstrap values >70 and were reproduced in a Bayesian phylogenetic analysis of a smaller dataset (Fig. S2) (SI Results).

**Figure 1.**
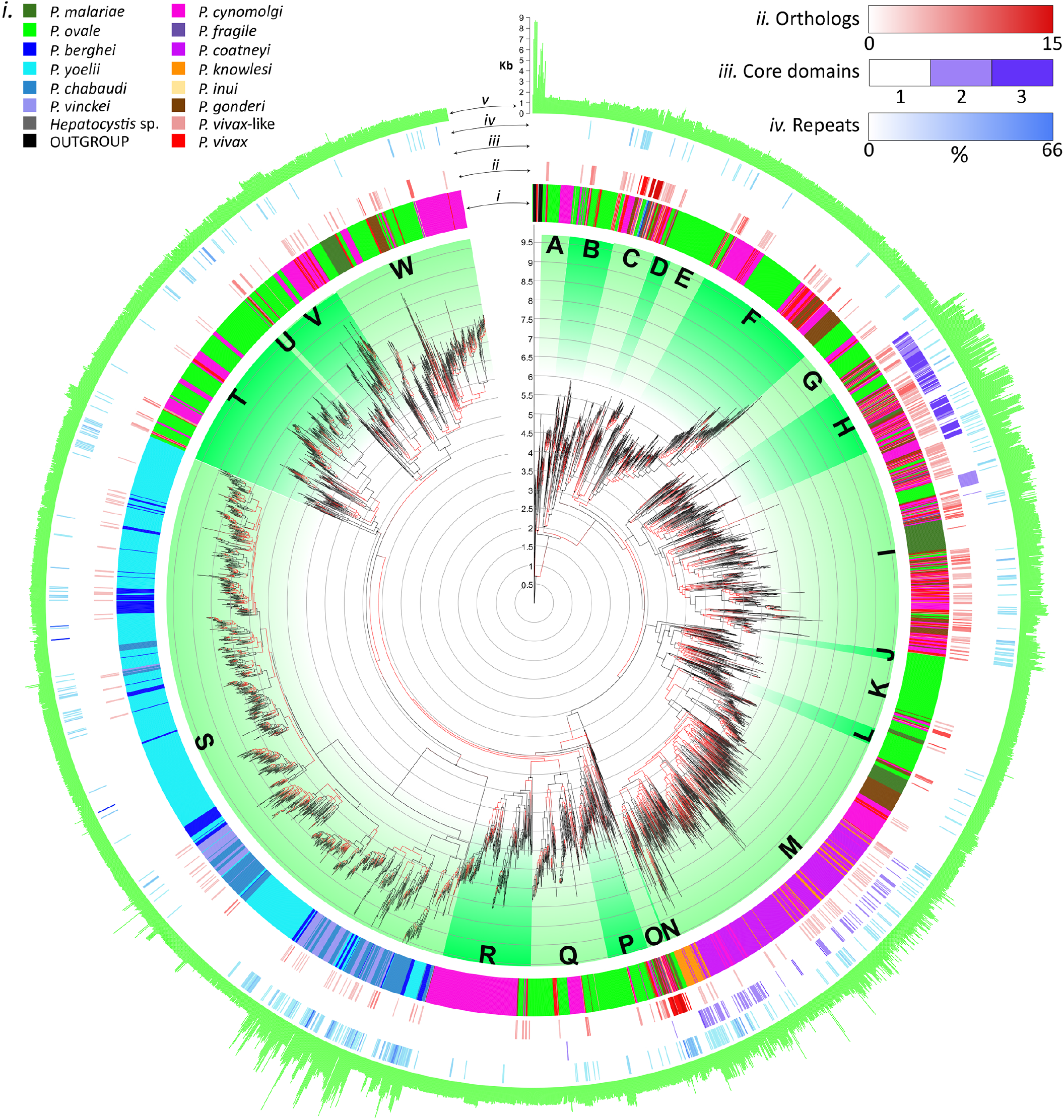
Maximum likelihood phylogeny of *Plasmodium* interspersed repeat (*pir*) protein sequences. The phylogram shown at centre was generated using IQTREE from an 837-character alignment of 4236 PIR protein sequences taken from 15 *Plasmodium* or *Hepatocystis* species genomes, supplemented with 17 SURFIN protein sequences, which are designated as the outgroup. A JTT+F+G model of amino acid substitution was applied with a Gamma shape (alpha) value of 3.16. Phylogenetic distance is indicated by a vertical scale. The phylogeny is subdivided into major clades thata are indicated by green segments and labelled as subfamilies A-W. Edge shading denotes branch robustness; black and red shading indicates edges that are subtended by nodes with bootstrap values >70 and <70 respectively. The phylogeny is surrounded by five tracks, from inside to outside: *i)* parasite species; *ii)* orthology (i.e. where a sequence belongs to an orthologous clade after reconciliation analysis (see text), the number of sequences in the clade is given); *iii)* number of conserved protein cores; *iv)* percentage of protein sequence comprising amino acid repeats; *v)* total protein length in amino acids. More intense shading represents higher numbers for orthology, cores and repeats respectively.

Species-specific clades are commonplace, indicating that gene duplication is frequent and that within-species paralogy is a dominant feature in *pir* evolution (see first track in Fig. 1). However, most subfamilies contain *pir* genes from multiple species (except *pirE* and *pirJ*, which are P. *ovale*-specific), indicating that some ancestral orthology is maintained in different *Plasmodium* species against a background of frequent gene duplications. For example, the high number of *pir* sequences in P. *ovale* and *P. cynomolgi* results from both the maintenance of ancestral subfamilies, coupled with substantial, species-specific gene duplications in these subfamilies. So, while *pirK* and *pirR* are found in diverse *Plasmodium* species, they are usually not abundant, however, *pirK* (102 paralogs) has expanded in P. *ovale* and *pirR* (156 paralogs) has expanded in *P. cynomolgi* in a species-specific manner.

Examining the predicted tertiary structures of *pir* subfamilies reveals that their early divergence coincided with structural differentiation, with each subfamily showing modifications of different regions of the ancestral protein (Fig. S3). For instance, *pirS* and *pirU* have evolved longer linker regions between the core and transmembrane domains (i.e. the *‘*distal variable*’* domain in Fig. S1), while *pirM* and *pirO* have expanded the region following the transmembrane domain (i.e. the *‘*proximal*’* domain). While protein length is relatively well conserved, with most *pir* genes encoding a protein of approximately 450 amino acids, longer *pir* genes, encoding proteins of >1,200 amino acids, have evolved on multiple occasions in several subfamilies. The *pir*S genes are divided into two clades, comprising short and long forms (Otto *et al*, 2014). Long PIRS proteins evolved through the expansion of amino acid repeats in the distal variable domain. In contrast, some PIRI proteins have evolved greater length through the expansion of repeats within the proximal domain, whereas long PIRH proteins have evolved through the duplication of the distal conserved domain, resulting in proteins that have multiple core domains. Finally, long *pirM* sequences have evolved through a combination of both core duplication and repeat expansion. Given this degree of structural difference between PIR proteins, the *pir* subfamilies defined here are highly likely to be functionally non-redundant and retained across the phylogeny because they have distinct roles.

By identifying *pir* subfamilies that are present in both rodent and primate genomes and thus derived from the common ancestor, we have established a universal phylogenetic systematics that standardizes comparisons of *pir* from known species and can be used to identify *pir* genes in new genomes and transcriptomes. As a proof of principle, we examined the transcriptomes of two lemur malaria parasites (Larsen *et al*, 2016) (Fig. S4, Data files 4 and 5). These parasites comprise a malaria lineage distinct from published *Plasmodium* genomes, yet possess *pir* genes that fit into our systematics, including the highly conserved *pirC1* gene (SI Results, Data file 6).

### The highly conserved *pir* is not the *‘*founder*’* of the *pir* family

Sequences belonging to the highly conserved gene (i.e. virD and its orthologs in other species) (Neafsey *et al*, 2012; Frech & Chen, 2013; Jackson, 2016; Little *et al*, 2021) form a clade (the *‘pirC1* clade*’*) within the *pir*C subfamily. All species considered here are represented in the *pirC1* clade; including all four *Hepatocystis pir* sequences. However, neither the *pirC1* clade sensu stricto, nor the wider *pirC* subfamily, can be the founder or ancestral clade from which all other *pir* genes have originated because *pirC* is not the sister clade to all other *pir* – the *pirA* and *pir*B subfamilies branch closer to the root. Constraining the phylogeny to make *pirC* the most basal branch results in a significantly less likely tree, which can be rejected (AU test, p=0.00032). Although the *pirC1* clade represents the only *pir* gene that is universally present, there are other instances where orthology is retained across multiple species. Another cluster of orthologs occurs in the *pirO* subfamily, which is among the rarest and most gene-poor subfamilies, but contains orthologs from all primate (though not rodent) malaria parasites, even in *P. fragile* and *P. inui* that have otherwise retained very few *pir* genes.

### The *pir* gene repertoire has varied extensively during *Plasmodium* evolution

Most *pir* genes (82.6%) are within-species paralogs, whose closest relati*ves* are near-identical copies in the same genome, while the remaining genes (17.3%) are orthologs (or paraphyletic to orthologs) that are most akin to *pir* genes in other species. To identify these conserved orthologs, we reconciled the *pir* phylogenetic tree with the *Plasmodium* species phylogenetic tree (Fig. 2). By inferring gene duplication and loss events, reconciliation must explain the observed distributions of genes in genomes, for instance, how most *pir* subfamilies contain genes from different *Plasmodium* species, and how rodent malaria parasite genomes, which are dominated by *pir*S, can also possess the distantly related *pir*C.

**Figure 2.**
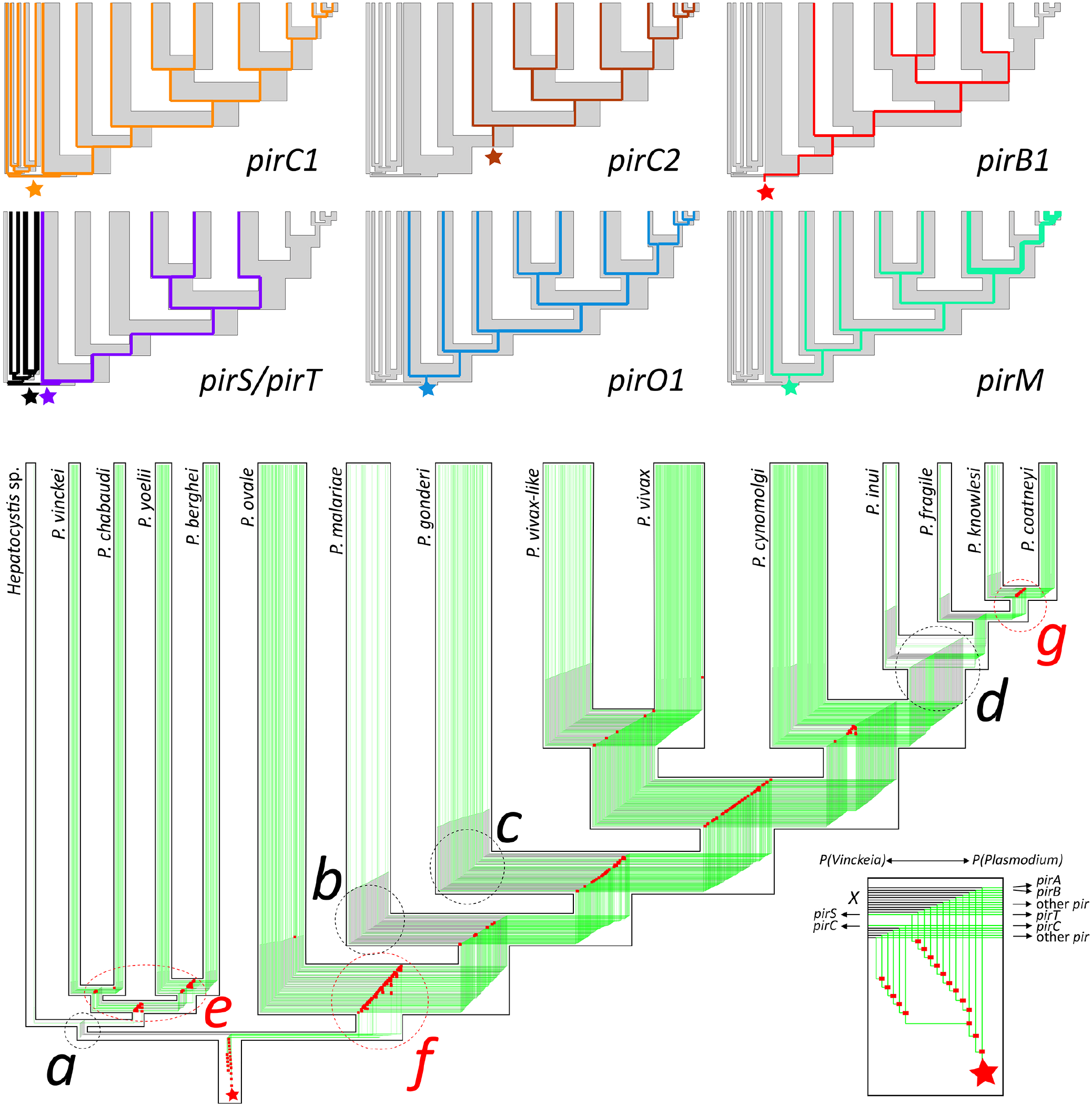
Reconciliation of *pir* and *Plasmodium* phylogenies. The *pir* gene tree was reconciled with the known relationships for the 15 species concerned here using TREERECS. The main output, showing the many genetic lineages fitted within the species tree, is shown in green. The common ancestor of all *pir* genes is shown by a red star. Gene duplications are indicated by red rectangles. Where deletion of a gene is inferred, this is shown with a black branch. For comparison, the gene trees of six specific *pir* clades are shown at top. The reconciled tree identifies points in species phylogeny that coincide with substantial reduction in *pir* diversity (**a-d**), coinciding with the origins of rodent malaria species (*Vinckeia* subgenus) and *Hepatocystis, P. malariae*, and the ancestor of *P. inuii, P. knowlesi, P. coatneyi* and *P. fragile* respectively. It also identifies moments of substantial expansion in *pir* diversity (**e-g**), coinciding, respectively, with diversification of rodent malaria species, origin of primate malaria species (*Plasmodium* subgenus) and diversification of the clade including *P. knowlesi* and *P. coatneyi*. (Inset) Expanded view of the *pir* gene tree root, showing basal duplication events that created the major *pir* subfamilies in the ancestor of *Vinckeia* and *Plasmodium* subgenera, (most of which were subsequently lost from *Vinckeia*).

The reconciled tree shows that a series of gene duplications in the genome of the common ancestor must be inferred to explain the contemporary distribution of *pir* genes (red nodes, Fig. 2 inset). The different subfamilies must have originated early in the common ancestor of *P(Vinckeia*) and *P(Plasmodium*); thereafter, the *pir* repertoire has fluctuated in both subgenera (Fig. 2). There have been phases of gene loss (labels a-d in Fig. 2), especially in the *P(Vinckeia*) subgenus, only the *pir*C and *pir*S subfamilies have been retained in this subgenus. Substantial gene loss has also occurred on the branches leading to *P. malariae*, and to *P. fragile*/*P. inui* /*P. knowlesi*/*P. coatneyi*. In the latter case, *pir* genes were almost entirely lost from the genomes of these species. Closer inspection of the *P. inuii* genome reveals numerous *pir ‘*gene relicts*’* that show evidence for this history of gene loss (SI Results).

In addition to gene loss, there have also been phases of gene duplication during *pir* evolution (labels e-g in Fig. 2), for example leading to rodent and primate malaria subgenera and in the ancestor of *P. knowlesi* and *P. coatneyi*. The expansion of *pir*S began in the common ancestor of rodent malaria genomes (but after the separation from *Hepatocystis*) and has continued as *P(Vinckeia*) species have diversified (Fig. 2 *‘*e*’*). It represents a secondary radiation following the deletion of most other *pir* subfamilies in *P(Vinckeia*), shortly after its separation from *P(Plasmodium*) (Fig. 2 *‘*a*’*). Although most primate malaria species possess *pirM* genes, this subfamily contributes the majority of *pir* genes in *P. knowlesi* and *P. coatneyi* genomes (Fig. 2 *‘*g*’*). This too represents a secondary radiation that has expanded the *pir* repertoire following its almost complete loss (Fig. 2 *‘*d*’*). The anomalous genomic positions of *pirM* gene loci in *P. knowlesi* and *P. coatneyi*, i.e. in chromosomal core regions instead of the subtelomeric positions characteristic of all other species, is consistent with their secondary evolution (Chien *et al*, 2016; Pain *et al*, 2008). Secondary radiation restored the mixture of *‘*short*’* and *‘*long*’* form *pir* genes that would have disappeared following periods of gene loss; both *pir*S and *pirM* subfamilies have evolved this length variation independently.

Changes to either the gene or species tree would alter these observations. Therefore, we assessed the effect of systematic error in the *pir* gene tree on the results of phylogenetic reconciliation by estimating and reconciling bootstrapped topologies. This did not change the timing of *pir* expansion and contraction phases described or the membership of orthologous clades (Fig. S5, Data File 7).

Hence, the *pir* gene content of contemporary *Plasmodium* species has arisen from three different trajectories of *pir* evolution: i) retention and expansion of all ancestral *pir* subfamilies (i.e. in P. *ovale*, P. cynomolgi, P. vivax), ii) loss of ancestral subfamilies followed by secondary radiation (i.e. in *P. malariae, P. knowlesi*/*P. coatneyi*, rodent malaria species), and iii) loss of ancestral subfamilies without replacement (i.e. in *P. inui, P. fragile, Hepatocystis*).

### Chromosomal synteny of conserved *pir* orthologs

Although species-specific gene loss and duplication has caused *pir* gene repertoire to fluctuate extensively during *Plasmodium* evolution, we noted above that several subfamilies include clusters of orthologs. Phylogenetic reconciliation identified 62 conserved orthologs across the tree, which are listed in Table S2. The *pirC1* genes was already documented but other genes displaying similar long-term orthology in primate malaria species include *pirC2* (e.g. PVX_110822), *pirC3* (e.g. PVX_115475) *pir*H4 (e.g. PVX_086893) and *pirO1* (e.g. PVX_096940). The latter is a highly conserved locus, present in all primate malaria parasites. In several species the *pirO1* locus consists of a tandem *pir* gene duplication and orthology is maintained among the tandem copies, suggesting that they are functionally differentiated (Fig. S6).

Although most *pir* genes are arranged in irregular arrays within the subtelomeric regions of chromosomes (Cunningham *et al*, 2010), some of the most widely conserved orthologs are present in conserved syntenic locations in different species. Of the 62 conserved *pir* genes, 17 (including the *pirC1, pirC2* and *pirO1* genes) are located in conserved loci closest to the internal core region of the chromosome, approximately at the boundary between the core and subtelomeric regions (Fig. S7, Table S3). Positional conservation of these select genes extends to *P. knowlesi* and *P. coatneyi*, in which, as noted above, *pir* loci have been substantially remodelled during a secondary radiation, and for *P. inui* and *P. fragile*, despite the extreme *pir* gene loss in these species. Not all conserved *pir* genes fit this pattern; some are located in subtelomeric regions distal from the core (e.g. *pirW1*).

### Amino acid replacement in *pir* gene orthologs is less frequent than in within-species paralogs

The phylogenetic and genomic position of the *pir* orthologs (Table 2), as described above, suggest that they have evolved differently from most other *pir* genes, which are typically within-species paralogs. We therefore examined the relative importance of mutation and selection in orthologous and paralogous *pir* genes. We first created codon alignments for orthologous *pir* clades where a test was possible (i.e. at least four genes; n = 20; Table S4), as well as for within-species paralogs of the same subfamily. CodeML (Yang, 2007) to compare models of codon evolution for orthologous and paralogous alignments and estimate the *D*_*n*_*D*_*s*_ ratio (ω) per codon. For *pirC1*, this analysis showed that an evolutionary model including positive selection was rejected, and 120/320 codons were estimated to be under negative selection (Fig. 3, Data file 8). By contrast, most codons in alignments of related *pir*C paralogs from four different species had a slightly negative ω (or occasionally positive), consistent with nearly-neutral evolution. Similar analysis of other subfamilies confirmed that this is a general pattern among conserved orthologs. Models including positive selection were rejected for 14/20 alignments of orthologs but preferred for 33/52 alignments of paralogs (Table S4). This pattern is most apparent in Fig. S8 (Data file 9), where the ω value across 14/20 alignments can be seen to be consistently lower for orthologs (green lines) relative to paralogs (black lines). Therefore, *pir* gene orthologs are rarely under positive selection, and more commonly negatively selected, in contrast to related *pir* gene paralogs.

**Figure 3.**
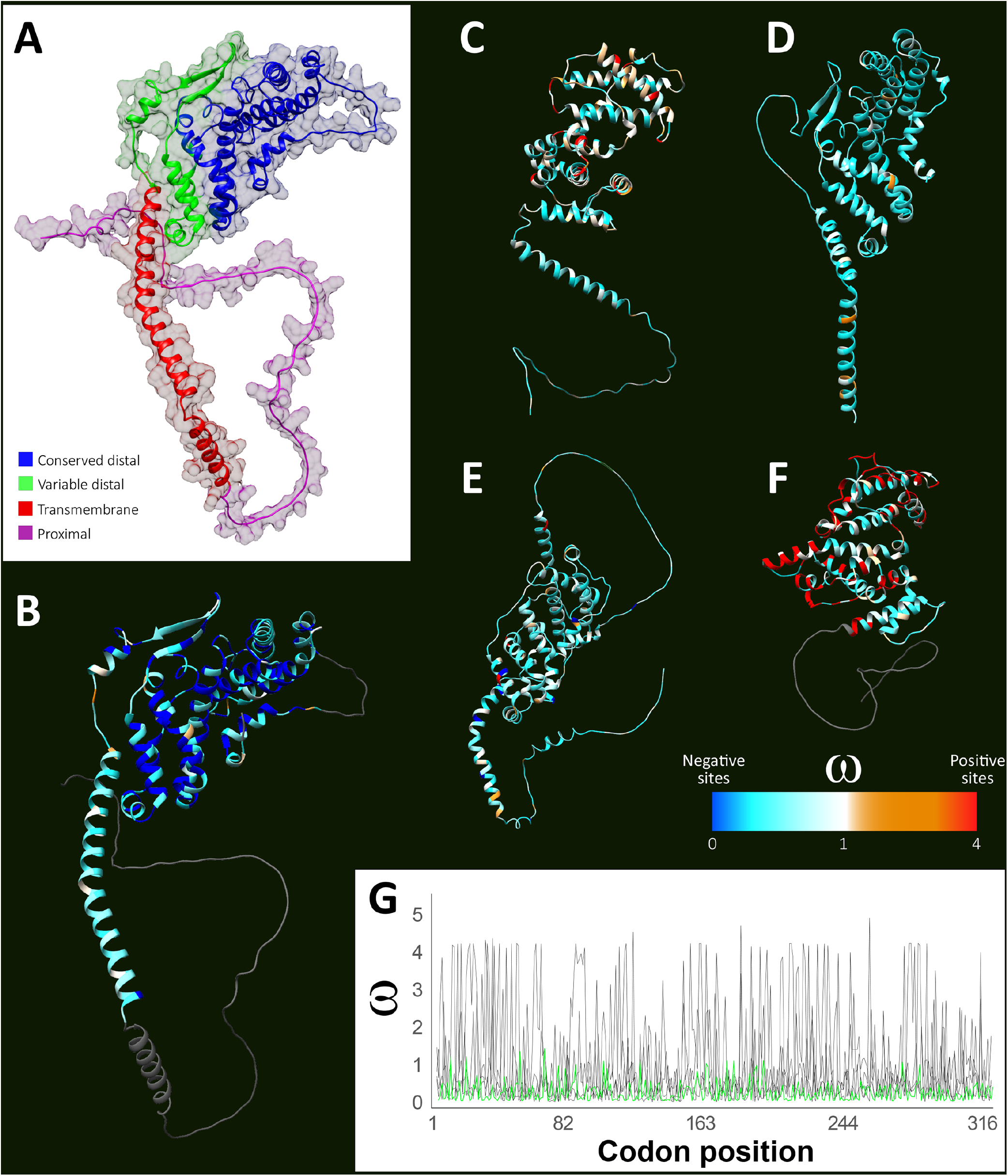
Estimation of selection across the PIRC1 protein structure compared to related paralogs. **a)** The protein structure of *pirC1*, as inferred by Alphafold (AF-K6V314); shading indicates four distinct regions of the sequence alignment (Fig. S1). **b)** *D*_*n*_*/D*_*s*_, the ratio of synonymous to non-synonymous amino acid substitutions, (ω) was estimated using codeML (Yang, 2007) for a codon alignment of *pirC1* ortholog sequences taken from 15 species. Individual residues in the protein model are shaded by the ω value of their corresponding codons. Sites showing significant negative selection, as determined by the Bayes em*pir*ical Bayes method of codeML (Yang *et al*, 2005), are shown in dark blue. Regions not included in the sequence alignment are shaded dark grey. c-f) For comparison, selection across four alignments of *pir*C within-species paralogs, each applied to their Alphafold predicted structures, are shown for *P. cynomolgi* (AF-K6V3I4), *Hepatocystis* (AF-A0A653GW70), P. *ovale* (AF-A0A1D3JEM5) and *P. vivax* (AF-A5KDQ9) respectively. Sites showing significant positive selection are shown in dark red. g*) D*_*n*_*/D*_*s*_ (ω) ratio values for *pirC1* codon sequences (green line) compared with the four groups of *pir*C within-species paralogs (black lines).

In summary, a select group of *pir* genes in multiple subfamilies is conserved in *Plasmodium* species (even in genomes where most *pir* genes are lost), have a syntenic genomic location (often at the core-subtelomeric boundary), and often evolve significantly slower than related, species-specific *pir* genes. Accordingly, when we mapped *pir* transcript abundance from published blood-stage transcriptomes of *P. vivax* (Zhu *et al*, 2016; Gural *et al*, 2018; Vivax Sporozoite Consortium *et al*, 2019) and *P. cynomolgi* (Cubi *et al*, 2017) to the *pir* phylogeny (Fig. S9), the conserved orthologs *pirC1, pirC2* and *pirO1* provided the most abundant transcripts across multiple experiments. Furthermore, single cell transcriptomic data in the Malaria Cell Atlas show that *pirC1* is expressed by all cells at specific points in the intraerythrocytic cycle of *P. knowlesi* (very early ring stage) and *P. berghei* (predominantly late schizont) (Fig. S10) (Howick *et al*, 2019). These characteristics indicate that *pirC1*, and other conserved orthologs, perform essential functions distinct from the majority of *pir* genes.

### Evidence for an essential role of *P. berghei* and *P. chabaudi* PIRC1 for optimal growth and localization of *P. berghei* PIRC1 in the parasitophorous vacuole

The *pirC1* gene is among 18 of the 68 *pir* genes that had a significant negative effect on the growth and multiplication of *P. knowlesi* blood stages when disrupted by genome-wide transposon mutagenesis (Elsworth *et al*, 2025; SI Results). To determine whether *pirC1* orthologs have a conserved, essential function in blood stages, we in*ves*tigated the function of *pirC1* in three genetically tractable *Plasmodium* species: the rodent malaria species *P. berghei* and *P. chabaudi* and the primate malaria species *P. knowlesi*. First, we attempted to delete the *pirC1* genes in *P. berghei* and *P. chabaudi* parasites using standard methods of gene deletion (Fig. S11A and Fig 12A-C). Using two different approaches (in 8 experiments total) no *P. berghei* parasites with a deleted *pirC1* gene (PBANKA_0100500) could be selected, strongly suggesting that *pirC1* is essential for *P. berghei* blood-stage growth (Fig. S11A). In contrast, a *P. chabaudi* mutant lacking *pirC1* (PCHAS_0101200) was obtained (Fig. S12). However, this *pirC1* mutant has a severe growth defect as shown by a longer patency period (approximately three days) after infection of mice and a significantly lower parasitaemia (p<0.01) during the acute phase (Fig. 4A). Observation of *P. chabaudi* blood stages in Giemsa-stained thin blood films revealed no obvious change in morphology of the *P. chabaudi* mutant (Fig. 4B, see additional images in Fig. S13).

**Figure 4.**
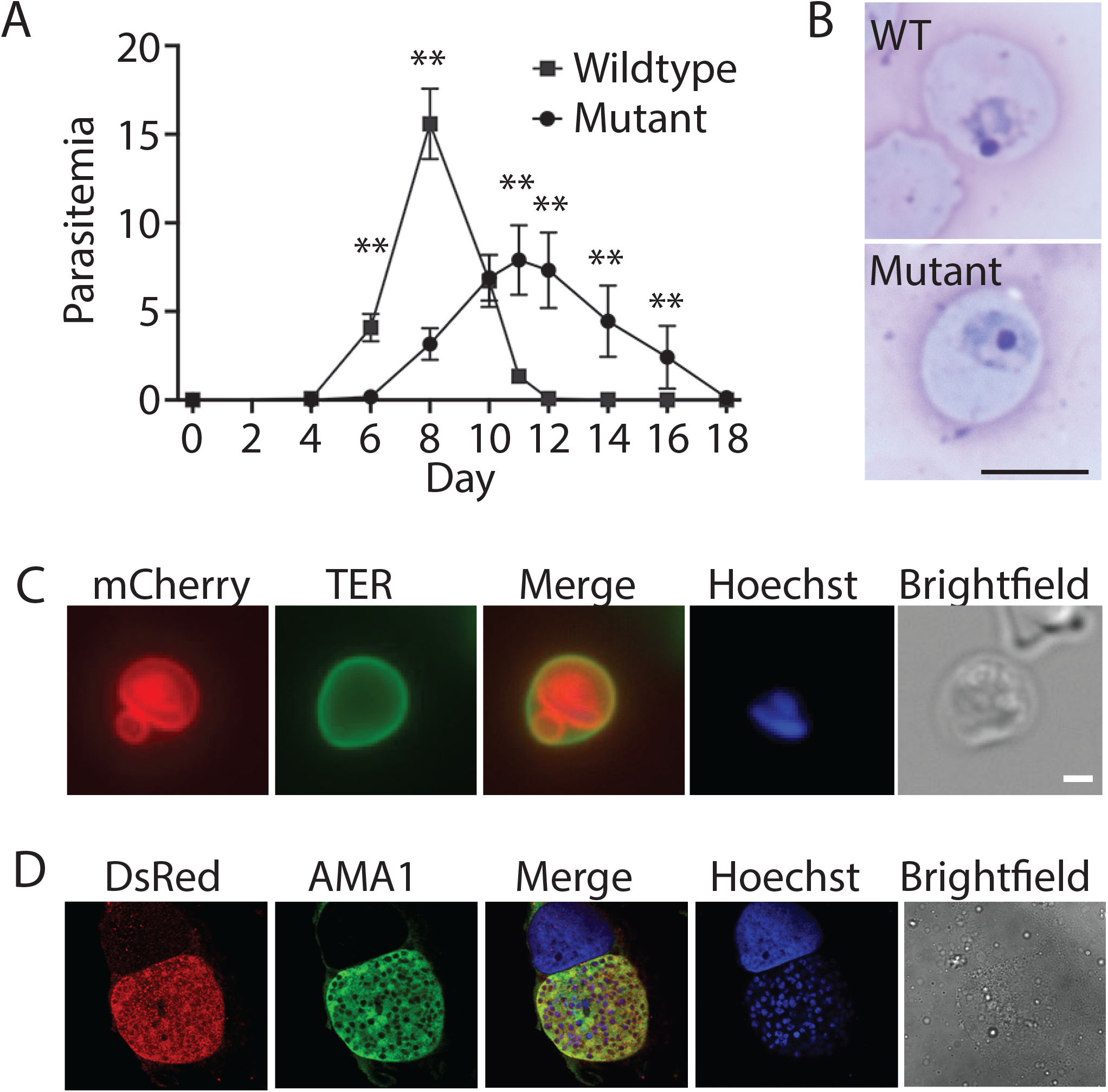
Growth of murine parasites lacking PIRC1. **a)** Replication of *Plasmodium chabaudi* control transgenic line *PcASluc*_*230p*_ (Cunningham *et al*, 2017) and *ΔPCHAS_0101200* parasites in C57BL/6 mice (n=8). Parasitaemia was monitored at 2-day intervals by enumeration of parasites on Giemsa-stained thin blood films. Note that the mutant displayed a longer patency and a lower maximal parasitemia; **p<0.01 (Mann-Whitney). **b)** Giemsa-stained thin smears of P. *chabaudi*-infected erythrocytes expressing *pirC1* (top) or lacking *pirC1* (bottom). See Fig. S13 for additional images. Bar indicates 5 μm. **c)** Live imaging of the localization of *P. berghei* PIRC1-mCherry in an erythrocyte infected with a trophozoite-stage parasite. The cells were stained with TER to visualize the erythrocyte membrane and Hoechst 33342 to visualize the parasites DNA – the lack of overlap of mCherry and TER indicates that little to no PIRC1-mCherry is exported. For images of additional intraerythrocytic stages, see Fig. S14. Bar equals 1 μm. **d)** Localization of PIRC1-mCherry *P. berghei*-infected hepatocytes. The parasites were visualized by staining with anti-AMA1 antibodies and PIRC1-mCherry was visualized using anti-mCherry antibodies. Host and parasite DNA was visualized by staining with Hoechst 33342.

These results indicate that PIRC is important for the optimal growth of *P. berghei* and P. *chabaudi*. This is the first direct demonstration of an essential role for a PIR protein.

To in*ves*tigate the possible function of PIRC1, we determined the cellular location of *P. berghei* PIRC1. For this, a mutant that expresses a C-terminal mCherry-tagged PIRC1 was generated (Fig. S11B-D). In blood stages the PIRC1-mCherry fusion protein was detected in a peripheral pattern around the parasite, indicative of a parasitophorous vacuole (PV) or parasitophorous vacuole membrane (PVM) location (Fig. 4C, see Fig. S14 for additional images) and was also detected in compartments extending into the infected erythrocytes in structures that resembled the tubo*ves*icular network (TVN), which is connected to the PV. mCherry fluorescence was also present in the cytoplasm of the blood stage parasites, but the PIRC1-mCherry fusion was not detected in the RBC cytoplasm or RBC membrane, as indicated by the lack of overlap with TER staining of the RBC membrane (Fig. 4C). The localization in the PV/PVM of PirC1 in *P. berghei* agrees with results of a proximity biotinylation study that identified PIRC1 as a PV protein (Schnider *et al*, 2018). PIRC1-mCherry was not detected in *P. berghei* mosquito stages but was detected in liver stages (Fig 4D), indicating that PIRC1 is produced by *P. berghei* during this stage.

### *P. knowlesi* PIRC1 is essential for growth

Since the results of the genome-wide transposon mutagenesis study in *P. knowlesi* indicated that lack of PIRC1 affected blood stage-growth (Elsworth *et al*, 2025; Oberstaller *et al*, 2025) and the PIRC1 orthologs have important role in blood stages of the rodent malaria parasites, we analysed the function by generating inducible gene-deletion mutants in *P. knowlesi*. Two *loxP* sites were introduced in the *P. knowlesi* gene (PKNH_1149300) in a parasite line that expresses diCre (Fig. S12D) (Collins *et al*, 2013; Mohring *et al*, 2020). Addition of rapamycin to cultures of blood stages of these *pirC1*-*loxP* parasites induces recombination between the *loxP* sites, thereby deleting most of the *pirC1* gene sequence (Fig 12E). Unexpectedly, after rapamycin treatment, the parasitaemia increased after 24 hours (the first developmental cycle), similar to the DMSO (vehicle)-treated cultures (Fig 5A). However, the parasitaemia in the rapamycin-treated cultures did not increase in subsequent cycles, remaining constant the following four cycles. In contrast, in DMSO-treated cultures, the parasitaemia increased to over 10% in subsequent cycles (Fig. 5A). Therefore, the lack of PIRC1 affects the growth of blood stages. The increase in parasitaemia of *P. knowlesi* lacking *pirC1* between the first and second developmental cycle is probably due to *pirC1* expression early in the intraerythrocytic cycle (in ring forms) and hence the protein is already present at the time of rapamycin treatment of the cultures that results in gene excision. Indeed, data from the Malaria Cell Atlas indicates that *P. knowlesi pirC1* is expressed at the very early ring stage (Fig. S10), whereas the *P. berghei pirC1* gene is expressed in the schizont stage, consistent with storage of the protein in merozoites with subsequent release after invasion (Howick *et al*, 2019). Futher supporting the requirement for the protein soon after merozoite invasion of erythrocytes is the finding that *pirC1* is highly expressed in liver merozoites (Little *et al*, 2021).

**Figure 5.**
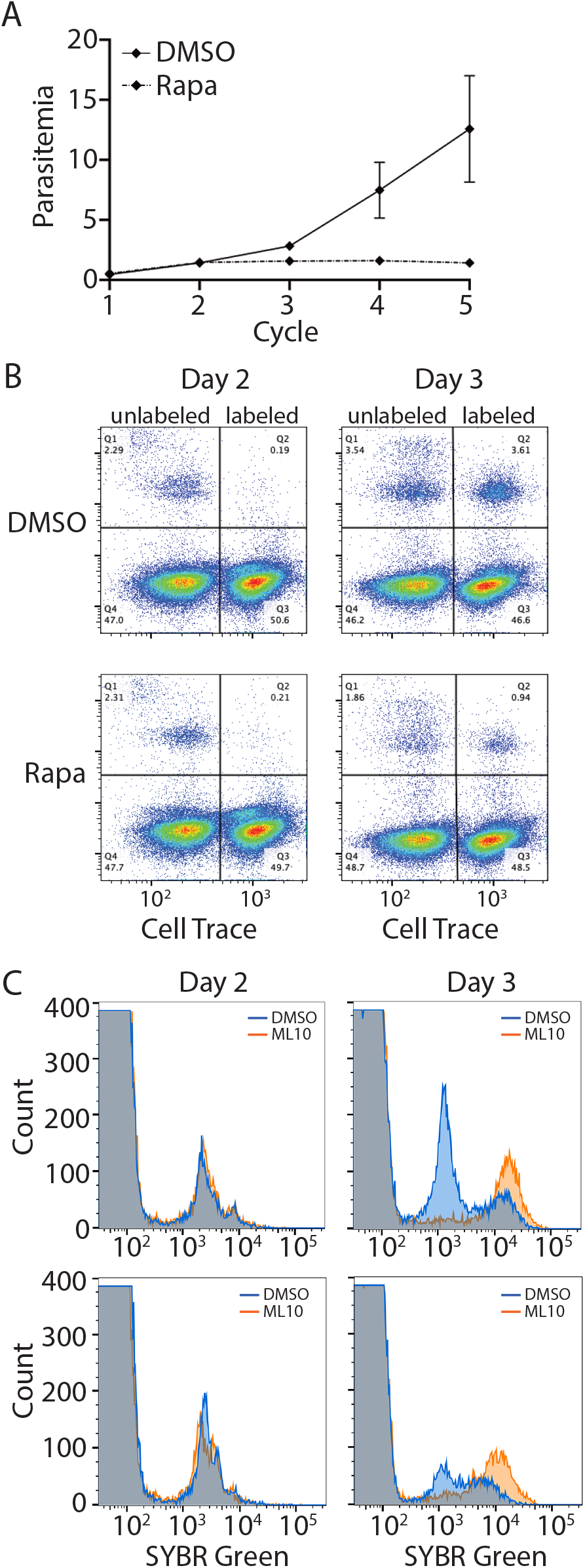
Growth of *Plasmodium knowlesi pirC1* mutant. **a)** Replication of *P. knowlesi* wildtype parasites and parasites lacking PIRC1. Synchronized parasites were treated with DMSO or 100 nM rapamycin during cycle 1 and the parasitemia was determined in the subsequent cycles by staining an aliquot of the culture with SYBR Green followed by cytometry. **b)** Invasion of *P. knowlesi* into erythrocytes labeled with CellTrace. Synchronized parasites were treated with DMSO or rapamycin on day 1 and CellTrace-labeled erythrocytes were added on day 2 such that approximately half the erythrocytes were labeled. Parasitemia was determined with SYBR Green staining and cytometry in all cycles. Note the appearance of SYBR Green-positive CellTrace-labeled cells (upper right-hand quadrant) in both the wildtype and mutant culture on day 3. Note also the lack of increase in the parasitemia of the rapamycin-treated parasites. **c)** DNA content of parasites treated with DMSO or rapamycin in the presence or absence of ML10. Synchronized parasites were treated with DMSO or rapamycin on day 1 and ML10 or DMSO was added to the cultures on day 2. DNA content was determined using SYBR Green and cytometry on day 2 and day 3. Note the accumulation of parasites with a high DNA content in the ML10-treated cultures of the wildtype and mutant parasites.

The lack of further increase in parasitaemia after the first developmental cycle of rapamycin-treated blood stages of the *pirC1* mutant could indicate that either the parasites are in a state of stasis, comparable to the cessation of development in parasites lacking exogenous isoleucine (Babbitt *et al*, 2012) and in parasites unable to produce polyamines (Assaraf *et al*, 1986), or that the parasites develop in the RBC through the entire intraerythrocytic cycle but that after egress of merozoites on average only a single merozoite invades a new erythrocyte (with a basic reproduction rate R_0_=1). To differentiate between these possibilities, we determined whether mutant parasites can invade new (labelled) erythrocytes and tested whether egress inhibitors arrest parasites at a late schizont, multinucleate stage.

To determine whether parasites lacking PIRC1 invade new erythrocytes, synchronized cultures of the *pirC1*-*loxP* parasites were treated with either rapamycin or DMSO (day 1) and allowed to progress to the next cycle (day 2). Uninfected erythrocytes, fluorescently labelled with CellTrace that can be identified by flow-cytometry (Theron *et al*, 2010), were then added to the culture so that half of the erythrocytes in the culture were fluorescent. In these cultures mutant parasites were detected in labelled erythrocytes 24 hours after the addition of the labelled erythrocytes (day 3) (Fig. 5B), indicating that merozoites had been released and had entered new erythrocytes. The fraction of parasites lacking PIRC1 in labelled erythrocytes was lower compared to that in the culture of DMSO-treated parasites on the third day, possibly indicating that the developmental cycle of the parasites lacking PIRC1 is slightly longer than that of parasites that produce PIRC1 (Fig. 5B).

To analyse whether *pirC1*-*loxP* parasites can progress through the complete developmental cycle, rapamycin-treated cultures of *pirC1*-*loxP* parasites were treated midway in the second cycle, approximately 36 hours after rapamycin or DMSO treatment, with the cGMP-dependent protein kinase (protein G) inhibitor ML10 that blocks egress of merozoites from mature schizonts (Ressurreição *et al*, 2020; Baker *et al*, 2017). If parasites lacking PIRC1 are in stasis it is expected that DNA replication to produce the mature schizonts is absent and that the DNA content remains the same 24 hours later when parasites are har*ves*ted. However, in the cultures treated with ML10 a population of parasites is present at 24/48 hour after start of the cultures with a higher DNA content than the parasites in the starting culture, in both cultures of wild type and rapamycin-treated *pirC1*-*loxP* parasites. Hence, parasites lacking PIRC1 can develop from ring forms into schizonts and become arrested at the late-schizont stage, similar to DMSO-treated parasites, when treated with ML10 (Fig. 5C). In contrast, in the control DMSO-treated and rapamycin-treated cultures that had not been treated with ML10, the DNA content of the parasites was similar on day 3 to that of culture on day 2, indicating that the parasites had egressed and invaded fresh erythrocytes. Note that whereas the parasitaemia in the DMSO-treated culture increases from day 2 to day 3, the parasitaemia in the culture of parasites lacking PIRC1 does not increase appreciably, as shown above.

Combined, these results indicate that *P. knowlesi* parasites lacking PIRC1 can replicate, egress and invade erythrocytes but have a very low basic reproduction rate of R_0_=1 compared to parasites that produce PIRC1. When we analyzed unsynchronized blood-stage parasite cultures, parasites lacking *pirC1*-*loxP* were clearly smaller than parasites producing PIRC1 (Fig. 6A). In Giemsa-stained thin blood films the mutant parasites appeared more compact, often with poor separation between the blue methylene blue staining of the parasite cytosol and the purple eosin staining of the nucleus. To determine when during the intraerythrocytic cycle this phenotype becomes apparent, we analysed synchronized cultures that were treated with DMSO or rapamycin. The parasites appeared normal 8 hours after invasion but the rapamycin-treated parasites were more compact than the DMSO-treated parasites 16 hours after invasion. After 24 hours, DMSO-treated *pirC1*-*loxP* parasites had developed into the schizont stage with a morphology comparable to wild type schizonts, whereas the rapamycin-treated *pirC1*-*loxP* parasites were noticeably smaller and appeared to contain fewer nuclei (Fig. 6B, see additional images in Fig. S15). Quantitation of the size of parasites at 24 hour after invasion confirmed that mutant parasites are significantly smaller than wild type parasites (p<0.001) (Fig. 6C) and quantitation of the DNA content of the DMSO-treated and rapamycin-treated *pirC1*-*loxP* schizonts arrested with ML10 (as shown in Fig. 5C), revealed that the DNA content in ML10-arrested parasites lacking PIRC1 is lower than that of DMSO-treated parasites (Fig. 6D). Together, these result show that PIRC1 is essential for the growth of *P. knowlesi*, even in when cultured in vitro, and functions early in the intraerythrocytic cycle.

**Figure 6.**
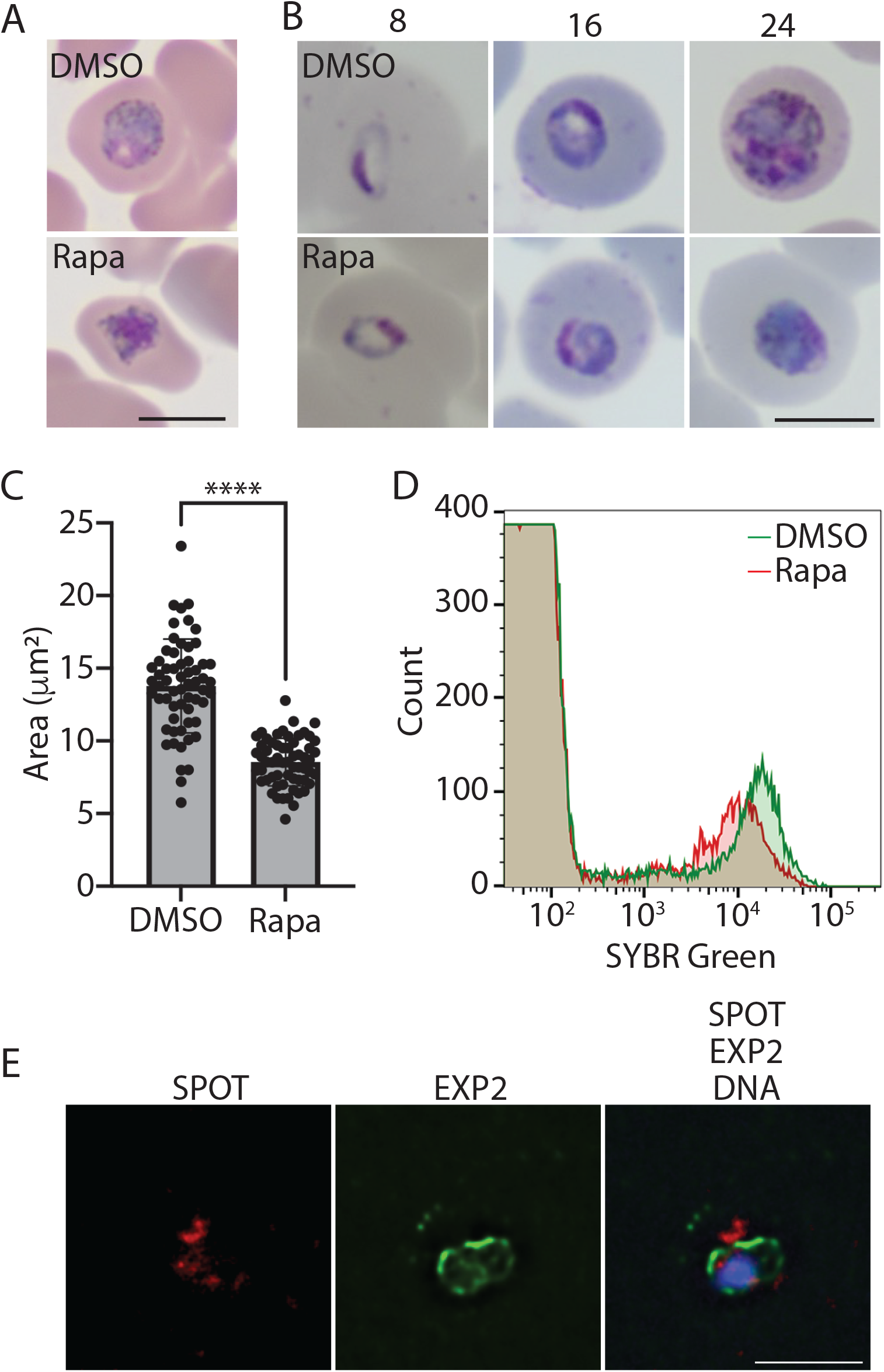
Phenotypic analysis of *P. knowlesi pirC1* mutant and localization of PIRC1. **a)** Giemsa-stained thin smears of erythrocytes infected with *P. knowlesi* parasites expressing (DMSO)) or lacking *pirC1* (Rapa). **b)** Development of *P. knowlesi* parasites with or lacking PirC1 after invasion. Synchronized parasites were treated with DMSO or rapamycin and synchronized again and blocked with ML10. Invasion of parasites was synchronized by removing ML10 and thin-film blood smears were produced at the indicated times. See Fig. S15 for additional images. **c)** Size of wildtype *P. knowlesi* parasites and parasites lacking PIRC1. Parasites were treated with DMSO or rapamycin and synchronized prior to passage to the subsequent intraerythrocytic cycle. Thin-film blood smears were stained with Giemsa and parasite size was determined by microscopy; ****p<0.001. **d)** Comparison of DNA content of the parasites producing or lacking PIRC1 shown in Fig. 5C. **e)** Localization of *P. knowlesi* PirC1 during the asexual intraerythrocytic cycle. PIRC1 was visualized using an anti-SPOT antibody and the PVM was visualized using an anti-EXP2 antiserum. Parasite DNA was visualized by staining with Hoechst.

### *P. knowlesi* PIRC1 is exported

Localization of PIRC1 by immunofluorescence microscopy of blood-stage parasites (using antibodies against the SPOT tag introduced at the C terminus of PirC1 (Fig. S12D)) revealed that the protein is exported into the RBC cytoplasm and located beyond the PV and PVM, as shown by costaining with antibodies against the PVM marker EXP2 (PF3D7_1471100) (Fig. 6E). This location in the RBC cytoplasm is in agreement with the observation that PIRC1 of *P. vivax* (i.e. VirD; PVX_113230) is transported into the cytoplasm of the RBC and associates with Maurer*’*s clefts in RBCs infected with P. *falciparum* that expresses *P. vivax* PIRC1 (Rehn *et al*, 2022). One location for PIRC1 may be the parasite exomembrane system in the cytoplasm of *P. knowlesi* infected RBC and the protein may function in its formation. We examined the exomembrane system in *P. chabaudi* and *P. knowlesi* parasites lacking PIRC1 and although PIRC1 may be associated with the exomembrane system, absence of PIRC1 does not affect its formation (Fig. S16).

## DISCUSSION

In this study we provide a comprehensive phylogeny of *pir* genes, showing that within-species paralogy is the dominant feature in *pir* evolution and provide evidence that at least on *pir* gene, *pirC1*, has an essential function. We identified 23 *pir* subfamilies, many of which contain orthologous *pir* genes that are conserved in multiple species, indicating that, despite a background of frequent gene duplications, some ancestral *pir* genes are maintained over long periods. To explain this, early origins of the different subfamilies in the ancestor of rodent and primate malarias must be posited, prior to separation of *P(Vinckeia*) and *P(Plasmodium*) but after their separation from *P(Laverania*). Thereafter, the *pir* repertoire has fluctuated as rodent and primate malaria parasites have diversified; these fluctuations have involved both *pir* gene loss and duplication, such that the *pir* repertoires of contemporary *Plasmodium* species have arisen from three different evolutionary trajectories – retention and expansion of all ancestral *pir* subfamilies, loss of ancestral subfamilies followed by secondary radiation, and loss of ancestral subfamilies without replacement.

The dominance of gene duplication in this family is expected given that previous phylogenies have contained species-specific clades or clusters (Otto *et al*, 2014; Jackson, 2016). Assuming that ancestral genomes were dominated by within-species paralogs, as they are now, most paralogs must be short-lived, not persisting long enough to leave orthologs. Gene conversion is likely to provide the mechanism for this rapid gene turnover. A genomic comparison of *P. chabaudi* subspecies showed that *pir* genes are created and deleted by gene conversion events at subtelomeric loci (Lin *et al*, 2018). This is likely to be true in other species; a cursory comparison of the genomes of P. brasilianum and *P. malariae*, (which are over 98% identical and probably conspecific (Bajic *et al*, 2022)), reveals similar gene conversion events at a conserved subtelomeric locus (Fig. S17). Indeed, the arrangement of *pir* genes in tandem arrays close to the telomeres of chromosomes will promote such non-homologous crossing-over events (Corcoran *et al*, 1988; Balzano *et al*, 2021), explaining why *pir* copy number fluctuates (Little *et al*, 2025) and why sequences commonly lack orthology within *Plasmodium* species. In *P. vivax*, for example, 62% of *pir* genes in a Peruvian Amazon strain (PvPAM) have no reciprocal *pir* sequence matches in a Papua Indonesia (PvP01) strain (De Meulenaere *et al*, 2023). We hypothesise that the relatively recent origins of within-species paralogs, and ease with which they are deleted, indicate that they are functionally redundant. This is supported by the rarity of strong purifying selection acting on within-species *pir* paralogs (Fig. S9) in favour of neutral or adaptive evolution.

However, against this background of rapid gene turnover, we have identified a select group of *pir* orthologs that are structurally differentiated and are often maintained in diverse species. We propose that these particular *pir* genes are functionally non-redundant. Previously, long-term conservation had only been reported for the virD gene of *P. vivax*, leading to its interpretation as the *‘*ancestor*’* or *‘*founder*’* of all other *pir* genes (Neafsey *et al*, 2012; Frech & Chen, 2013; Little *et al*, 2021). Jackson defined such a gene family member, which robustly forms the most basal branch of a phylogeny, closest to the outgroup, as a *‘*protolog*’* (Jackson, 2016); Group I Serine Repeat Antigen genes (i.e. PfSERA8) (Arisue *et al*, 2007) and the PHISTc subfamily (Sargeant *et al*, 2006) are examples in *Plasmodium* parasites. The *pirC1* gene, equivalent to the clade comprising virD and its orthologs, is indeed unique among *pir* genes as being universally conserved in all *Plasmodium* species that possess *pir* genes. However, when the *pir* phylogeny is rooted with *surfin*s, the *pirC1* clade is not the most basal branch, nor the sister clade to all other *pir* genes. Therefore, *pirC1* is not the protolog from which all other *pir* genes have originated. Instead, it is one of several genes that originated in the common ancestor of rodent and primate malarias and have persisted since.

This apparent essentiality of *pirC1* throughout *Plasmodium* evolution agrees with our observations of severe growth defects in blood stages of *P. berghei, P. chabaudi* and *P. knowlesi*. In these species the deletion of the *pirC1* gene had a variable effect. Whereas *P. berghei* blood stages lacking *pirC1* could not be selected, *P. chabaudi* and *P. knowlesi pirC1* gene deletion mutants were viable, albeit with a severe growth defect. The growth rate of blood stages of *P. knowlesi pirC1* mutants in vitro was approximately one parasite per cycle while wild type parasites have a mean growth rate in vitro of 2.5-3 parasites per cycle in human blood (Moon *et al*, 2013). We cannot discount the possibility that these phenotypic differences among *pirC1* mutants of different species may reflect the presence of closely related *pir* genes (e.g. *pirC2* in *P. knowlesi*) in some species that might partially complement the lack of *pirC1*.

In addition to revealing the essential role for PIRC1, this study provides insight into its cellular function. The requirement for *pirC1* for optimal growth of *P. knowlesi* in vitro of blood stages cultures rules out a function in immune evasion or immunomodulation. Based on the increase in *P. vivax pirC1* expression in late schizonts, a role for *pirC1* in invasion was suggested (Neafsey *et al*, 2012). However, as *P. knowlesi* parasites lacking *pirC1* can progress to the next cycle in fresh RBCs, *pirC1* is unlikely to be essential for invasion. The decreased proliferation, the reduced level of DNA at late blood stages and the aberrant morphology of *P. knowlesi* blood stages lacking *pirC1* are more consistent with a metabolic defect. As PIRC1 is from the parasite itself and is present at the parasite-host interface, either in the PV/PVM or the exomembrane system, PIRC1 may play a role in the acquisition of nutrients from the RBC, either directly or by stabilizing or activating an uptake or transport mechanism. Interestingly, *Hepatocystis* sp. only possess *pirC1*, while its lifecycle lacks an asexual cycle in RBC and produces gametocytes directly from liver stage merozoites (Aunin *et al*, 2020). We show that PIRC1 is present during the *P. berghei* liver stage (Fig. 4D), while transcriptomic analyses have shown that the *pirC1* gene is transcribed during the late liver stage, asexual stage and gametocyte stages of multiple species (Little *et al*, 2021). Thus, although parasites lacking *pirC1* have a severe phenotype in asexual blood stages, its presence in *Hepatocystis* parasites and other life cycle stages of *Plasmodium* prasites suggests that *pirC1* function relates to a basic parasite-host interaction, potentially the uptake of nutrients, at multiple life cycle stages.

Other orthologous *pir* genes (i.e. *pirC2, pirC3, pirB1, pirP1* and *pirO1*) display similar evolutionary profiles to *pirC1* and, by extension, warrant further in*ves*tigation into essential roles in cell growth and development. Conversely, the function of the majority of species-specific, paralogous *pir* genes may be different. Having proposed that rapid evolutionary turnover reflects functional redundancy, we suggest that the number and diversity of *pir* sequences is functionally important per se rather than a specific, conserved protein structure as with *pirC1*. Thus, *pir* genes may have a role in immune modulation or evasion (Cunningham *et al*, 2010), cytoadherence (Bernabeu *et al*, 2012; Carvalho *et al*, 2010) and in modulating virulence (Harrison *et al*, 2020; Brugat *et al*, 2017). Ultimately, the phylodynamics of *pir* genes are not uniform and strongly support multiple, distinct functions.

## CONCLUSION

Most *pir* genes are relatively recently evolved paralogs that can be individually or entirely lost under some circumstances, suggesting that they are functionally redundant. Remarkably for one of the largest *Plasmodium* gene families, there are evidently conditions during evolution in which *pir* genes are almost dispensible. Thus, the phylodynamics of most *pir* genes do not indicate long-term essentiality; rather, the profusion and rapid evolution of *pir* repertoires suggest a flexible and adaptive response to changing host-parasite interactions. In stark contrast, a minority of *pir* genes are ancient, long-term orthologs with evolutionary dynamics that do not indicate exposure to immune selection or rapid evolution. We have shown that the most highly conserved of these genes, *pirC1*, is essential for growth and localises to the host-parasite interface, suggesting a role in nutrient acquisition or host sensing. Several other *pir* genes (*pirC2, pirO1*) are likewise so important that they resist deletion during evolution, even when every other *pir* gene is lost, and these may be propitious targets for experimentation. The main lesson from the phylogeny when considering *pir* function is to distinguish old orthologs from young paralogs, since their evolutionary differences are very likely derived from fundamentally different roles.

## MATERIALS AND METHODS

### Sequence sampling and alignment

Amino acid sequences of proteins encoded by *pir* genes were recovered from published genome sequences for four rodent *Plasmodium* species (*P. berghei, P. chabaudi, P. yoelli, P. vinckei*), 10 primate *Plasmodium* species *(P. cynomolgi, P. vivax, P. vivax-like, P. malariae*, P. *ovale, P. inui, P. fragile*, P. gonderi, *P. knowlesi, P. coatneyi*) and one *Hepatocystis* species held in PlasmoDB (release 67) and/or GenBank. BLASTp was used to identified *pir* sequences among annotated proteins from each genome using recognised PIR proteins from *P. cynomolgi* and *P. berghei* as a query. With an initial alignment of the BLASTp matches generated using ClustalW (Thompson *et al*, 1994) a hidden Markov model (HMM) was generated using HMMER 3.3.2 (Eddy, 2011) that was used to search the predicted proteomes for more distant homologs. These searches did not examine pseudogenic sequences, which we initially excluded to produce an alignment that included full-length and intact gene copies. Table S1 records the number of annotated *pir* (including pseudogenes) in each genome to show how many *pir* sequences were omitted. Some pseudogenic sequences were subsequently added to the alignment when they occurred in conserved positions (i.e. they belonged to conserved orthologous clades). To provide an outgroup for comparative analysis, 17 *surfin* sequences from five species *(P. cynomolgi, P. vivax, P. ovale, P. falciparum, P. relictum*) were added. See SI Results for justification of their selection as outgroup. 4,232 *pir* and *surfin* amino acid sequences were aligned using ClustalW and manually adjusted in BioEdit (Hall, 1999) producing 839 characters. Many sequences contain complex and hypervariable repeat regions that are not well aligned; these were largely removed to leave an alignment of the conserved alpha helical and transmembrane domains (SI Results).

### Phylogenetic estimation

A Maximum Likelihood phylogeny was estimated using IQ-TREE 2.0.7 (Minh *et al*, 2020) on the CIPRES platform (Miller *et al*, 2010, 2015). Automated model selection identified JTT+F+G as the optimal amino acid substitution model. Tree robustness was evaluated with 1,000 ultrafast bootstraps. A Bayesian phylogeny was estimated using the same substitution model with BEAST v2.7.7 (Bouckaert *et al*, 2019). To allow tree parameters to reach convergence, a reduced data set of 142 sequences, comprising 4-10 chosen to reflect the diversity of each *pir* subfamily plus 17 outgroup sequences as before, was used in Bayesian analysis. The tree was estimated with four parallel Markov Chain Monte Carlo (MCMC) runs of one million generations, yielding 10,000 trees (and otherwise default settings). The resultant tree file was examined using Tracer (Rambaut *et al*, 2018) to ensure convergence of tree parameters based on sufficient expected sample size (ESS). The consensus tree was generated with a burn-in of 1,000 trees. The position of *pirC1* was evaluated using an approximately-unbiased (AU) test of tree topology (Shimodaira, 2002) and the reduced data set of 142 sequences described above. The optimal maximum likelihood tree was compared to constrained topologies in which either *pirC1* alone or the entire *pir*C subfamily were forced into the most basal position to test the hypothesis that *pirC1* represents the most basal branch of the *pir* phylogeny (i.e. the *‘*ancestor*’*). The phylogeny was annotated using iToL v7 (Letunic & Bork, 2024) with several properties: species, orthology (from reconciliation analysis), number of core domains (counted manually from alignment) and proportion of repeat sequence (estimated with XSTREAM (Newman & Cooper, 2007)).

## Phylogenetic reconciliation

Where gene trees disagree with species relationships, gene duplications and losses may be inferred to explain the differences, producing a reconciled tree. Phylogenetic reconciliation was carried out using TREERECS 1.2 [https://gitlab.inria.fr/Phylophile/Treerecs] and NOTUNG 2.9.1.5 (Chen *et al*, 2000) to reconcile the optimal *pir* tree with the *Plasmodium* species tree. The gene tree was produced by pruning the optimal Maximum Likelihood topology such that all species-specific clades were reduced to a single representative (n=730). This had the effect of removing all within-species paralogs, which was done to simplify the reconciled tree and focus attention on the origins (and losses) of between-species paralogs (i.e. *pir* subfamilies). In practice, this only affects very recent gene duplications that concern one species only. The tree was rooted with *pirA* sequences. The species tree was estimated using phylogenomics. Predicted proteomes for all 15 species concerned here plus five outgroups (*P. falciparum, P. gaboni, P. relictum, P. gallinaceum and Haemoproteus tartakovskyi*) were obtained from PlasmoDB and clustered using Orthofinder (Emms & Kelly, 2019). Orthogroups containing single-copy orthologs for at least 15/20 species were selected (n=2931) and concatenated using MISPhyl (Tan *et al*, 2013), then degapped using Gblocks 0.91b (Talavera & Castresana, 2007). A Maximum Likelihood phylogeny was estimated from the concatenated amino acid alignment using IQ-TREE2 with an LG+F+G model and 1000 ultrafast bootstraps. A partitioned model (LG+C20+F+G) accounting for rate differences between constituent sequences of the data set was applied but did not affect the result. A Bayesian phylogeny was estimated using the same substitution model with BEAST as described above.

The TREERECS output, which contains clades of all sizes, was filtered to select only non-redundant clades with orthologs from at least three species (given its closeness to *P. vivax, P. vivax-like* was not counted as a species for this purpose); in-paralogs from only one species were permitted. Larger clades containing multiple orthologs and smaller clades with fewer than three species represented were discarded, leaving only the smallest orthologous groupings with the most minimal paralogy. This identified *pir* genes conserved across species, which were labelled sequentially by subfamily and in order of decreasing size (i.e. most conserved first). To assess the effect of systematic error within the gene tree on the number of gene duplications and losses in the reconciled tree, alternative trees were generated. 100 bootstrapped trees were estimated using the pruned alignment described above and PHYLIP seqboot (https://phylipweb.github.io/phylip/doc/seqboot.html) followed by IQ-TREE2. These bootstrapped gene trees were each reconciled with the species tree (which was robust and accurate throughout) using NOTUNG 2.9.1.5, to estimate the number of duplication and loss events associated with each internal and terminal node of the species tree. Event numbers in the optimal gene tree were compared to the range observed among bootstrapped trees to assess the effect of uncertainty.

### Identification of *pir* genes from lemur malaria transcriptomes

Whole blood transcriptomes for two lemur species (*Propithecus diadema* and *Indra indra*) infected with malaria parasites were assembled from published RNA-seq data (PRJNA293089) (Pacheco *et al*, 2011) using SPAdes v4.0.0 (Prjibelski *et al*, 2020). Host reads were subtracted from the sequence data by mapping to the lemur genome with Hisat2 (Kim *et al*, 2019) and the remaining non-host reads were assembled. BLASTx was used to identify transcripts with a best hit to *Plasmodium* proteins, including PIRs. To determine the phylogenetic affinity of the lemur malaria parasites, putative lemur malaria amino acid sequences (N=735 and 580 for *P. diadema and I. indri* respectively) were clustered with predicted proteomes for the 15 species concerned here plus *P. falciparum* as an outgroup. 270 clusters of single-copy orthologs present in all genomes/transcriptomes (taking the longest transcript where several were available), were concatenated using MISPhyl and degapped using gBLOCKs, producing an alignment of 13260 characters. A phylogenomic tree was estimated using IQ-TREE2 and BEAST as above using an JTTDCMut+F+I+G4 model and rooted with *P. falciparum*. To determine the affinity of lemur malaria PIRs, these were added to the complete *pir* alignment and the phylogeny was re-estimated as before. To evaluate whether the subfamilies designated here could accommodate the *pir* sequences present in the lemur blood transcriptomes, HMMs for each subfamily were created with HMMER v3.3.2 and used to screen each translated transcript. The optimal subfamily (as determined by HMM score) and the observed phylogenetic placement of each transcript were then compared and were expected to coincide.

### Comparative genomics

To explore the genomic position of conserved, orthologous *pir* genes, tBLASTx result files were generated for pair-wise comparisons of chromosomal ends for all 14 chromosomes (i.e. from telomere to chromosomal core boundary) between species with adequate genome assemblies (i.e. *P. vivax, P. cynomolgi, P. gonderi, P. malariae, P. knowlesi, P. coatneyi* and *P. berghei*). Genome assemblies for *Hepatocystis* sp., P. *ovale, P. inui, P. fragile* and *P. vivax-like* do not include subtelomeres articulated with chromosomal cores and were not used. These comparisons were inspected using Artemis Comparison Tool (ACT) (Carver *et al*, 2005) to check the locations of orthologs, either predicted by tree topology or present in other species. A *‘*conserved*’* locus was recorded where orthologs were present in three or more of the seven species considered. If any species were found to have pseudogenes in such conserved positions, these were retrospectively added to the alignment and phylogeny. For *pirO1*, the genomic comparison in ACT was annotated with an accompanying phylogeny of the orthologs concerned (Fig. S5).

### Codon evolution

*D*_*n*_*/D*_*s*_, the ratio of synonymous to non-synonymous amino acid substitutions per site (ω) was estimated using codeML (Yang, 1997, 2007) for codon alignments of *pir* sequences and compared between conserved orthologs found in different species and within-species paralogs of the same subfamily. Clades identified by TREERECS were the guide for creating 20 alignments of orthologous *pir*. Unlike the main alignment, these codon alignments used the full-length gene sequences, including variable repeat regions where these were present, because closely related orthologs align along their full length (unlike the gene family as a whole). Ortholog alignments contained a minimum of four sequences from different species (typically from P. cynomolgi, *P. vivax, P. vivax-like*, P. gonderi or *P. malariae* where positional information could corroborate orthology), but possibly more if the gene was more widely conserved (e.g. *pirC1, pirC2* and *pirO1*). P. *ovale* orthologs were occasionally used if a strong argument could be made for orthology from the phylogeny alone (no positional information being available). Paralog alignments for comparison with each ortholog had at least four sequences from the same subfamily but potentially more depending on the species. Paralog alignments were created for multiple species if possible. To test the evidence for positive selection in each alignment, codeML estimated the likelihood of five site models for each alignment using the following settings: runmode = user tree, seqtype = codons, CodonFreq = F3X4, clock = no clock, model = one, NSsites = 0 1 2 7 8, icode = universal code, fix_omega = estimate, cleandata = no. Log-likelihood ratio tests (LRT) were used to compare likelihood values (L) for M1 (nearly neutral) vs. M2 (positive selection), and for M7 (beta) vs. M8 (beta with ω). Significantly greater likelihood for both M2 and M8 would indicate positive selection. Significance of the LRT statistic (i.e. 2ΔL) was inferred from the χ2 distribution with two degrees of freedom; thus, there is a significant difference between models where 2ΔL > 13.8 at p = 0.001. We also counted the number of negatively and positively selected codons for each alignment using the Bayes Em*pir*ical Bayes function (BEB) (Yang *et al*, 2005) in codeML. Negative selection was inferred for codons where the combined posterior probabilities for the first 3 ω classes > 0.95. Positive selection was inferred for codons where the combined posterior probabilities for the last 3 ω classes > 0.95.

### Experimental animals for *P. berghei* culture and reference *P. berghei* parasite line

Female OF1 mice (6–7 weeks; Charles River, NL) were used. All animal experiments were granted with a license by Competent Authority after an advise on the ethical evaluation by the Animal Experiments Committee Leiden (AVD1160020171625). All experiments were performed in accordance with the Dutch Experiments on Animals Act (Wod, 2014), the applicable legislation in the Netherlands, in accordance with the European guidelines (EU directive no. 2010/63/EU) regarding the protection of animals used for scientific purposes. All experiments were executed in a licensed establishment for the use of experimental animals (LUMC). Mice were housed in individually ventilated cages furnished with autoclaved aspen woodchip, fun tunnel, wood chew block and Nestlets at 21 ± 2°C under a 12:12 h light-dark cycle at a relative humidity of 55 ± 10%. To generate the PBANKA_0100500::mCherry and ΔPBANKA_0100500 parasites, the reference *P. berghei* ANKA WT (cl15cy1) line was used (Janse *et al*, 2006).

### Transmission of P. berghei

Mosquitoes from a colony of Anopheles stephensi (line Nijmegen SDA500) were used. Larval stages were reared in water trays (at 28⍰± ⍰1°C; 80% relative humidity). Adult females were transferred to incubators at 26v±⍰0.2°C (80% relative humidity) and were fed with 5% filter-sterilized glucose solution. For the transmission experiments, 3 to 5 day-old mosquitoes were used. Following infection, the *P. berghei* infected mosquitoes were maintained at 21°C at 80% relative humidity.

### Attempts to generate *P. berghei* parasites lacking expression of PBANKA_0100500

To replace the open reading frame of the conserved *pirC1* gene PBANKA_0100500, we created cloned the 5⍰ and 3⍰ flanking regions of the PBANKA_0100500 open reading frame on either side of a pyrimethamine-resistance selection cassette of pL0001 (Pasini *et al*, 2013) containing the *Toxoplasma gondii dhfr/ts*, generating pL2016. Primer pairs 7416-7417 and 7418-7419 were used to amplify the 5*’* region and 3*’* region, respectively and were cloned using Asp718 and HindIII (5*’* region) and EcoRV and XbaI (3*’* region). In addition, a construct was generated using the adapted *‘*Anchor-tagging*’* PCR-based method as described previously (Annoura *et al*, 2012). The two targeting fragments bir_0100500 were amplified as template using primer pairs 7481-7482 (5*’* target sequence) and 7483-7484 (3*’* target sequence). Transfection, selection and cloning of transfected parasites were performed as described (Janse *et al*, 2006). Transfected parasites were selected with pyrimethamine or WR99210 (de Koning-Ward *et al*, 2000; Janse *et al*, 2006). Correct integration of the DNA constructs was determined by diagnostic PCR and Southern blot analysis of chromosomes separated by pulse-field gel electrophoresis. Southern blots were hybridized with the 3*’*UTR dhfr/ts of *P. berghei* ANKA probe (Annoura *et al*, 2012) (PMID: 22342550). See Table S5 for the sequence of the primers.

### Generation and genotyping of parasites expressing PBANKA_0100500 tagged with mCherry

To generate transgenic parasites expressing a C-terminal-tagged mCherry tagged version of the PBANKA_0100500, construct pL1419 was modified as follows (Fonager *et al*, 2012). The *smac* targeting region was removed using SpeI and BamHI and replaced by a targeting region of the bir_0100500 that had been amplified using primers 7185 and 7186 to create plasmid pL1940. The inserted PCR fragment was sequenced after TOPO TA (Invitrogen) sub-cloning. Transfection of *P. berghei* parasites with linearized plasmid or PCR product, selection and cloning of transgenic and mutant parasite lines were performed as described (Janse *et al*, 2006).

### Analysis of expression of PBANKA_0100500::mCherry in blood and liver stages

For analysis of mCherry expression, tail blood of infected mice or infected erythrocytes were collected in PBS or complete RPMI-1640 and were examined by flow cytometry (see below) or fluorescence microscopy using a Leica DMR fluorescent microscope with standard GFP and Texas Red filters. Parasite nuclei were labelled by staining with Hoechst-33258 (Sigma-Aldrich, The Netherlands) and red blood cell surface membranes were stained with the anti-mouse TER-119-FITC labelled antibody (eBioscience, The Netherlands). Briefly, erythrocytes were stained with TER-119-FITC antibody (1:200) and Hoechst-33258 (2 µM) at room temperature for 30 minutes, cells were pelleted (400 x g, 2 minutes) and washed with 500 μL of RPMI-1640). For DNA visualization, Hoechst-33258 (2 µM) was added during the incubation with the TER antibody. Cells were pelleted (400 x g, 2 min) and suspended in RPMI-1640 medium.

The percentage of blood-stage parasites that expressed mCherry was determined by flow cytometric analysis of cultured blood stages. In brief, infected tail blood (10 µL) with a parasitemia between 1 and 3% was cultured overnight in 1 ml complete RPMI-1640 at 37°C under standard conditions for the culture of *P. berghei* blood stages (Fougère *et al*, 2016). Cultured blood samples were then collected and stained with Hoechst-33258 (2 µM; Sigma-Aldrich, The Netherlands) for 1 hour at 37°C in the dark and analysed using a FACScan (BD LSR II, Becton Dickinson, CA, USA) with filter 440/40 for Hoechst signals and filter 610/20 for mCherry fluorescence. For flow cytometry analysis the population of mature schizonts was selected based on their Hoechst-fluorescence intensity and the percentage of mCherry-expressing parasites was calculated by dividing the number of mCherry-positive schizonts by the total number of mature schizonts (Pasini *et al*, 2013).

Feeding of Anopheles stephensi mosquitoes was performed as described (Fougère *et al*, 2016). *P. berghei* PBANKA_0100500::mCherry sporozoites were isolated from salivary glands of infected A. stephensi mosquitoes 18-24 days after an infectious blood meal. The human hepatocyte carcinoma cell line Huh7 (JCRB0403, JCRB Cell Bank, Japan) was used for in vitro cultures of the liver stages. For immunofluorescence analysis of liver stages, 5×10^4^ sporozoites were added to a monolayer of Huh7 cells on coverslips in 24-well plates in RPMI-1640 supplemented with 10% (vol/vol) fetal bovine serum (FBS), 2% (vol/vol) penicillin-streptomycin and 1% (vol/vol) GlutaMAX (Invitrogen) and maintained at 37°C with 5% CO_2_. At 48 hours after infection, cells were fixed with 4% paraformaldehyde, permeabilized with 0.5% Triton-X 100 in PBS, blocked with 10% FBS in PBS, and subsequently stained with primary and secondary antibodies overnight at 4°C and for 1 hour, respectively. Primary antibodies used were anti-DsRed (632496, Takara Bio, Japan) and anti-AMAI 18G2 (kindly provided by Clemens Kocken, BPRC, The Netherlands). Secondary antibodies used were respectively anti-rabbit antibodies conjugated to Alexa Fluor 594 (A-21207, Invitrogen) and anti-rat antibodies conjugated to FITC (A18872, Thermo Fisher Scientific). Nuclei were stained with Hoechst-33342. Cells were mounted in Image-iT FX Signal Enhancer (Molecular Probes) and examined using a TCS SP8 Leica fluorescence microscope. Images analysis was done with the Leica LAS X software.

### Culture and mutagenesis of *Plasmodium chabaudi*

All experiments using *P. chabaudi* were performed in accordance with the UK Animals (Scientific Procedures) Act 1986 (PPL 70/8326 and PADD88D48) and were approved by The Francis Crick Institute Ethical Committee. V(D)J recombination activation gene RAG1 knockout (RAG1^−/−^) (Mombaerts *et al*, 1992) (Mombaerts *et al*., 1992) on a C57BL/6 background and C57BL/6 WT mice were obtained from the specific-pathogen-free (SPF) unit and subsequently conventionally housed with autoclaved cages, bedding and food at the Biological Research Facility (BRF) of the Francis Crick Institute. Experiments were performed with six-weeks to eight-weeks-old female mice under reverse light conditions (light 19:00-07:00 and dark 07:00-19:00), at 20-22°C.

A cryopreserved stock of a cloned line of *Plasmodium chabaudi chabaudi* AS, originally obtained from David Walliker, University of Edinburgh, UK, and subsequently passaged through mice by injection of infected red blood cells (iRBC), was the recipient line for transfection.

The PcAS *ΔPCHAS_0101200* construct was generated by replacement of the *P. chabaudi smac* region in the PcAS Δsmac construct described previously (Cunningham *et al*, 2017) with the *P. chabaudi PCHAS_0101200* targeting region (PCHAS_01_v3 50360-49650 and PCHAS_01_v3 52515-51810) (see Table S5). The resulting *ΔPCHAS_0101200* plasmid was digested with Kpn1/SacII digested was used to transfect the stock line as described previously (Spence *et al*, 2011) and *Plasmodium chabaudi* AS parasites in which the targeting DNA had integrated, and hence lacked expression of PCHAS_0101200, were selected with pyrimethamine (35 μg/ml) acidified drinking water. After passage into C57BL/6 mice, parasite genomic DNA was extracted from infected blood using a Zymogen DNA extraction kit according to manufacturer*’*s instructions. Integration into the *PCHAS_0101200* locus and presence or absence of the intact locus was verified by PCR using primers 2/3 and primers 1/3, respectively (Fig. S12 and Table S5).

RNA was extracted from the *ΔPCHAS_0101200* parasite line and wildtype parasites as described previously (Nahrendorf *et al*, 2015) and transcription of the *PCHAS_0101200* gene and the 18S housekeeping gene (PCHAS_0937240) was in*ves*tigated by QRT-PCR. *PCHAS_0101200* gene primers (Table S5) were designed using IDT Primer Quest software (www.idtdna.com), optimised and amplification efficiency assessed on serially diluted genomic DNA from wildtype parasites. Amplification was performed on a QuantStudio™ 3 System (45 cycles, TaqMan™ Fast Advanced Master Mix; Thermofisher) using primer and probe concentrations detailed in Table S5.

qRT-PCR amplification of the *ΔPCHAS_0101200* parasites and wild-type *P. chabaudi* AS parasites showed the absence of *PCHAS_0101200* transcription in the *ΔPCHAS_0101200* line after 45 amplification samples (CT value =45). Control parasites exhibited high expression of PCHAS_0101200, with CT values of 25-27. All samples expressed the 18S housekeeping (PCHAS_0937240) gene (CT values of 17-34 and 16-18) (Figure 2C and Table S5).

Six-week to eight-week old female C57BL/6 mice were infected intraperitoneally with 10^5^ infected red blood cells from *ΔPCHAS_0101200* parasite line (n=8) and a control transgenic line *PcASluc*_*230p*_ (described in (Cunningham *et al*, 2017) (n=8). Mice were maintained under drug selection throughout the infection as described above. Parasitaemia was monitored at 2-day intervals by enumeration of parasites on Giemsa-stained thin blood films.

### Culture and mutagenesis of *Plasmodium knowlesi*

*Plasmodium knowlesi* parasites were maintained at 37° C in an atmosphere of 96% N_2_, 1% O_2_, 3% CO_2_ in RPMI supplemented with 0.5% AlbuMax type II (Gibco), 50 μM hypoxanthine, 2 mM L-glutamine and 10% horse serum (cRPMI) and containing a hematocrit of 2-3%. Parasitemia was between 0.5-10% (Moon *et al*, 2013).

*P. knowlesi* parasites were synchronized on a Percoll cushion according to standard procedure (Ressurreição *et al*, 2022). Briefly, most of the medium was aspirated from the parasite culture and the erythrocytes were suspended in the remaining medium. This was gently layered on 2.5 ml of 65% Percoll in 1x RPMI, 3% sorbitol and centrifuged at 1200 x g for seven minutes with slow acceleration and brake. Late-stage parasites were transferred from the interface to a new 15 ml conical tube and 8 ml of cRPMI was added. This mixture was centrifuged at 1200 x g for four minutes with maximum acceleration and brake. The supernatant was aspirated and the parasites were suspended in 5 ml of cRPMI and transferred to a T75 flask containing 1 ml of packed erythrocytes. This culture was incubated at 37°C for 1-2 hours on a shaking platform. The appearance of ring-stage parasites in the culture was checked by Giemsa staining of thin-film smears. Remaining late-stage parasites were removed by a second floatation on a Percoll cushion, removing the infected erythrocytes at the interface and retaining the pelleted erythrocytes containing the newly invaded parasites.

The inducible *P. knowlesi* PKNH_1149300 mutant (*pirC1*-*loxP*) was produced by transfecting late-stage schizonts of the *P. knowlesi* PkdiCre line 7 (Hart *et al*, 2023) with the repair plasmid pBLD546 and the Cas9-guide RNA plasmid pBLD551 using the standard transfection protocol (Mohring *et al*, 2019). Transfectants were selected with 100 nM pyrimethamine for five days and resistant parasites were recovered after approximately 10 days. The parasites in the resulting culture were cloned by limiting dilution and integration of the repair plasmid in the isolated clones was verified by PCR using primer pairs CVO449-CVO365 (5*’*) and CVO433-CVO450 (3*’*) to amplify wildtype DNA and primer pairs CVO447-CVO365 (5*’*) and CVO433-CVO488 (3*’*) to amplify integrant DNA. This revealed that integration of pBLD546 had occurred using the 5*’* homology region and an 11 bp region in the region that was homologous to the wildtype sequence. To insert the second *loxP* site, the wildtype 3*’* region of PKNH_1149300 was amplified using primers FM1515 and FM1512, which also introduced sequence encoding the SPOT tag. The resulting fragment was amplified by PCR using primers FM1515 and FM1514 to produce a fragment that contains the *loxP* site. Sequence downstream of PKNH_1149300 was amplified in FM1513 and FM1000. This fragment was fused to the fragment with PKHN_1149300 3*’* region, SPOT tag and *loxP* sequence by overlapping PCR. The resulting fragment was ligated with the pCR ZeroBlunt plasmid to produce plasmid pBLD700. Parasites containing the 5*’* region of pBLD546 were transfected with this repair plasmid in conjunction with Cas9-gRNA plasmid pL10kir. The transfected parasites were selected as described above and cloned by using the plaque assay (Thomas *et al*, 2016). Correct integration of pBLD700 was verified using primers CVO681-FM1514 to detect wildtype sequence and CVO681-CVO683 to detect integrant sequence. Excision of the kir gene after treatment with rapamycin was determined using primer pair CVO446-FM1594. See Table S5 for primer sequences.

pBLD546 was produced by inserting a recodonized version of PKNH_1149300 compassing bp 789-1954 of the native gene. In this recodonized version, the first intron from the PkSERA2 gene in which a *loxP* site had been inserted was placed between bp 2591 and 2592 and the 5*’* region of the second native intron was replaced with the 5*’* region of the second intron from PkSERA5. This fragment was ligated to pCR Zero Blunt to produce pBLD546. pBLD551 was produced by annealing the oligos CVO370 and CVO380 and cloning the resulting double-stranded fragment into pDC2-Cas9-hDHFRyFCU (Knuepfer *et al*, 2017) that had been linearized with BbsI.

Gene deletion was induced as described previously (Collins *et al*, 2013; Knuepfer *et al*, 2017). Briefly, two 4-ml aliquots of parasite culture containing recently invaded parasites were placed in 15 ml conical tubes. In one tube, 4 ul of DMSO was added (for a dilution of 1:1000) and to the other tube 4 ul of 100 uM rapamycin in DMSO was added (for a final concentration of 100 nM). The cultures were incubated at 37°C for 45 to 60 minutes, after which the parasites were pelleted by centrifugation at 1200 x g for four minutes. The supernatant was aspirated and the parasites were suspended in 4 ml cRPMI, transferred to a T25 flask and incubated at 37°C.

### Growth assays and cytometry

Proliferation of parasites was determined as described previously (Fréville *et al*, 2024). Briefly, the parasite culture was treated with either rapamycin to induce deletion of the PKNH_1149300 or DMSO as described above. The parasitaemia was then adjusted to 0.1%, 0.5% or 1.0% in triplicate and 1 ml was transferred to a 24-well plate. A 50 ul aliquot was removed, mixed with 50 ul 2x fixative (8% paraformaldehyde, 0.2% glutaraldehyde) and stored at 4°C. Additional 50 ul samples were collected at 24-hour intervals. When all samples had been collected, the cells were prepared for cytometry as follows. Parasites were pelleted at 2000 x g and suspended in 100 ul PBS containing 1X SYBR Green and incubated for 15-60 minutes, after which the parasites were pelleted at 2000 x g and resuspended in 1 ml PBS. A 200 ul aliquot of each sample was transferred to a well of a 96-well plate and the fluorescence within the erythrocytes was determined using an Attune cytometer outfitted with a Cytkick using the following settings: forward scatter 125 V, side scatter 350 V, blue laser (BL1) 530:30 280V. The experiment was set up in triplicate, with a minimum of three biological replicates.

### Labelling of erythrocytes with Cell Trace and analysis of transition to next cycle

To determine the invasion of parasites lacking PIRC1 into fresh erythrocytes, infected erythrocytes were mixed with erythrocytes stained using the CellTrace Far Red Cell Proliferation Kit (Molecular Probes) (Theron *et al*, 2010). Erythrocytes (500 μl packed cells) were pelleted and washed three times with 3 ml Hanks*’* Balanced Salt Solution (HBSS). The cells were then suspended in 1 ml HBSS containing 1 μl of CellTrace Far Red dye (reconstituted as per manufacturer*’*s instruction) and incubated at 37°C for one hour. The erythrocytes were subsequently pelleted and suspended in 15 ml cRPMI, to produce a hematocrit of 3%.

To prepare parasites for the experiment, *pirC1*-*loxP* parasites were treated with 100 nM rapamycin or an equivalent volume of DMSO as described above. The parasitemia of the treated culture was adjusted to 1% and 1 ml of the mixture was placed in the well of a 24-well plate. A sample for cytometry was removed at this point. After 24 hours, the medium in the well was removed and the infected erythrocytes were suspended in 1 ml of fresh medium. One half of the suspension (500 μl) was placed in a well of a 24-well plate and 500 μl of Cell Trace-labelled erythrocytes was added. A aliquot for cytometry was removed at this point and on the two subsequent days. These aliquots were prepared for cytometry as described above and analysed using an Attune cytometer outfitted with a Cytkick (Thermo Fisher) with the following settings: forward scatter 125 V, side scatter 350 V, blue laser (BL1) 530:30 280V, red laser (RL1) 670:30 nm 250 V.

### Analysis of transition between cycles using ML10

To determine whether *P. knowlesi* parasites lacking cKIR are arrested at a late developmental stage, *pirC1*-*loxP* mutant parasites and A1.H.1 wildtype parasites were treated with DMSO or rapamycin as described above. After 24 hours, ML10 or an equivalent volume of DMSO was added to the culture to a final concentration of 150 nM and a small aliquot of the culture was removed for cytometry. The subsequent day, another small aliquot of the culture was removed to determine the effect of ML10. The parasites were prepared for cytometry and the DNA content was determined using staining with SYBR-Green as described above.

### Imaging of SPOT-tagged *P. knowlesi* PIRC1

Parasites were prepared for immunofluorescence as described previously (Fréville *et al*, 2024). Thin-smear blood films were air dried and fixed with 4% paraformaldehyde in PBS for ten minutes. After washing with PBS, the cells were permeabilized with 0.1% Tween-20 in PBS and subsequently blocked with 3% BSA in PBS. The cells were incubated in the presence of anti-SPOT antibody (1:1000; ChromoTek) and anti-EXP2 antibody for one hour at room temperature, washed and incubated in the presence of goat anti-mouse Alexa 488 and anti-rabbit Alexa-568 secondary antibodies (Invitrogen). The samples were washed, covered with VectaShield (Vector) and a glass coverslip. The parasites were imaged on a Nikon Ti-E inverted microscope with Hamamatsu ORCA-Flash 4.0 Camera and Piezo stage driven by NIS elements version 5.3 software. Images were futher processed with FIJI and Abobe Photoshop and figures were produced with Abode Illustrator.

### Labeling erythrocytes with fluorescent ceramide

Parasites were stained with fluorescent ceramide as described previously (Grüring & Spielmann, 2012). Briefly, a 100 μl aliquot of parasite culture was washed twice with RPMI without AlbuMax. The cells were subsequently pelleted, suspended in RPMI containing 14 μM C5-Bodiy-ceramide complexed to BSA (Fisher Scientific) and incubated at 37°C for 30 to 60 minutes. The cells were then washed twice with RPMI without AlbuMax, suspended in 400 μl HBSS and 200 μl of this was loaded into a well of a 6-well Ibidi poly-L-lysine coated μ-Slide (Thistle Scientific) slide and imaged on a Nikon Ti-E inverted microscope as described above.

### Imaging of Giemsa-stained parasites

Size of the parasites in Giemsa-stained thin films was determined as described before (Fréville *et al*, 2024). The slides were imaged on an Olympus BX51 microscope equipped with an 100x oil objective and an Olympus SC30 camera controlled by CellSense software. The size of the parasites was determined by measuring the longest diameter in each parasite using the line selection tool and set measurement function in FIJI/ImageJ software.

## Supporting information

Date File 3 for Figure 1

Data File 7 for Figure S5

Data file 8 for Figure 3

Data file 9 for Figure S8

Supplementary results

## Acknowledgments

We thank Clemens Kocken (BPRC, The Netherlands) for sharing the anti-AMA1 antibody 18G2 and Paul Gilson (University of Melbourne, Australia) for sharing the anti-EXP2 antiserum. This work was supported by a UKRI Medical Research Council Career Development Award (MR/R008485/1) and Wellcome Trust Institutional Strategic Support Fund (204928/Z/16/Z) to CvO. R.W.M, F.M and G.P were supported by a UKRI Medical Research Career Development Award (MR/M021157/1) and Wellcome Trust Discovery Award (225254/Z/22/Z). JL and DC were supported by the Francis Crick Institute, which recei*ves* its funding from the UK Medical Research Council, Cancer Research UK, and the Wellcome Trust, UK (FC001101). JL was a Wellcome Senior In*ves*tigator (WT 104777/Z/14/Z). This work was furthermore supported by Leiden University Medical Center internal funds. The funders had no role in the design, analysis or reporting of these studies. The authors acknowledge the facilities and the scientific and technical assistance of the LSHTM Wolfson Cell Biology Facility, with specific thanks to Liz McCarthy and Chris Chiu.

## Author contributions

A.P.J.,, D.A.C., T.S.L., R.W.M., J.L., C.J.J., B.M.D.F-F and C.v.O. designed research; A.P.J., D.A.C. L.L., N.M.C.d.O., S.C.C-M., G.P., F.M, A.K.R. and C.v.O. performed research; F.M. contributed new reagents/analytic tools; A.P.J., D.A.C, T.S.L., J.L., C.J.J., B.M.D.F-F. and C.v.O. analysed data; A.P.J., R.W.M., J.L., C.J.J. and C.v.O. acquired funding and provided resources and A.P.J. and C.v.O. wrote the paper.

