## Supplementary Figures 1-18 for "Essential function reflected in the phylodynamics of a multigene family – the *pir* genes of malaria parasites"

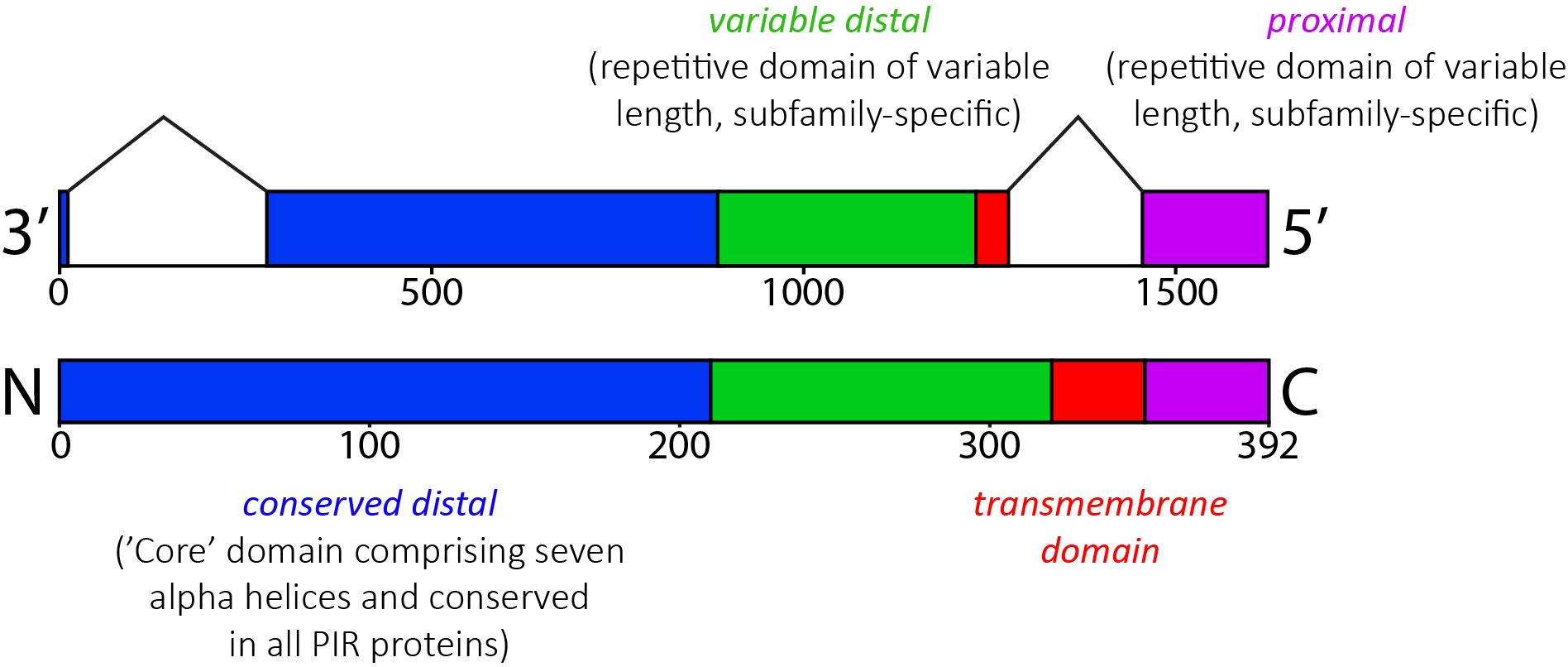

**Supplementary Figure 1. Cartoon of *Plasmodium* interspersed repeat (*pir*) gene and protein sequence structures.** The structure shown represents the *pirC1* gene in *P. vivax* (PVX_113230), which consists of three exons and two introns (top). The predicted protein sequence (bottom) is divided into four domains based on the global sequence alignment (see SI Results). A core domain consisting typically of the first seven alpha helices of the PIR protein (‘distal conserved’ – light blue). Another alpha helix is predicted to be a transmembrane domain (‘transmembrane domain’ – red). The light blue and red domains are joined by an often-repetitive region of variable length (‘distal variable’ - green). Following the transmembrane domain at the C-terminal of the PIR protein there is another, often repetitive region of variable length (‘proximal’ – purple). In situ, the protein is positioned across a membrane (potentially of parasitophorous vacuole membrane or erythrocyte plasma membrane). The ‘distal conserved’ and ‘distal variable’ domains are predicted to be orientated beyond the membrane, pointing away from the parasite. The ‘proximal’ domain is predicted to be orientated before the membrane, closest to the parasite.

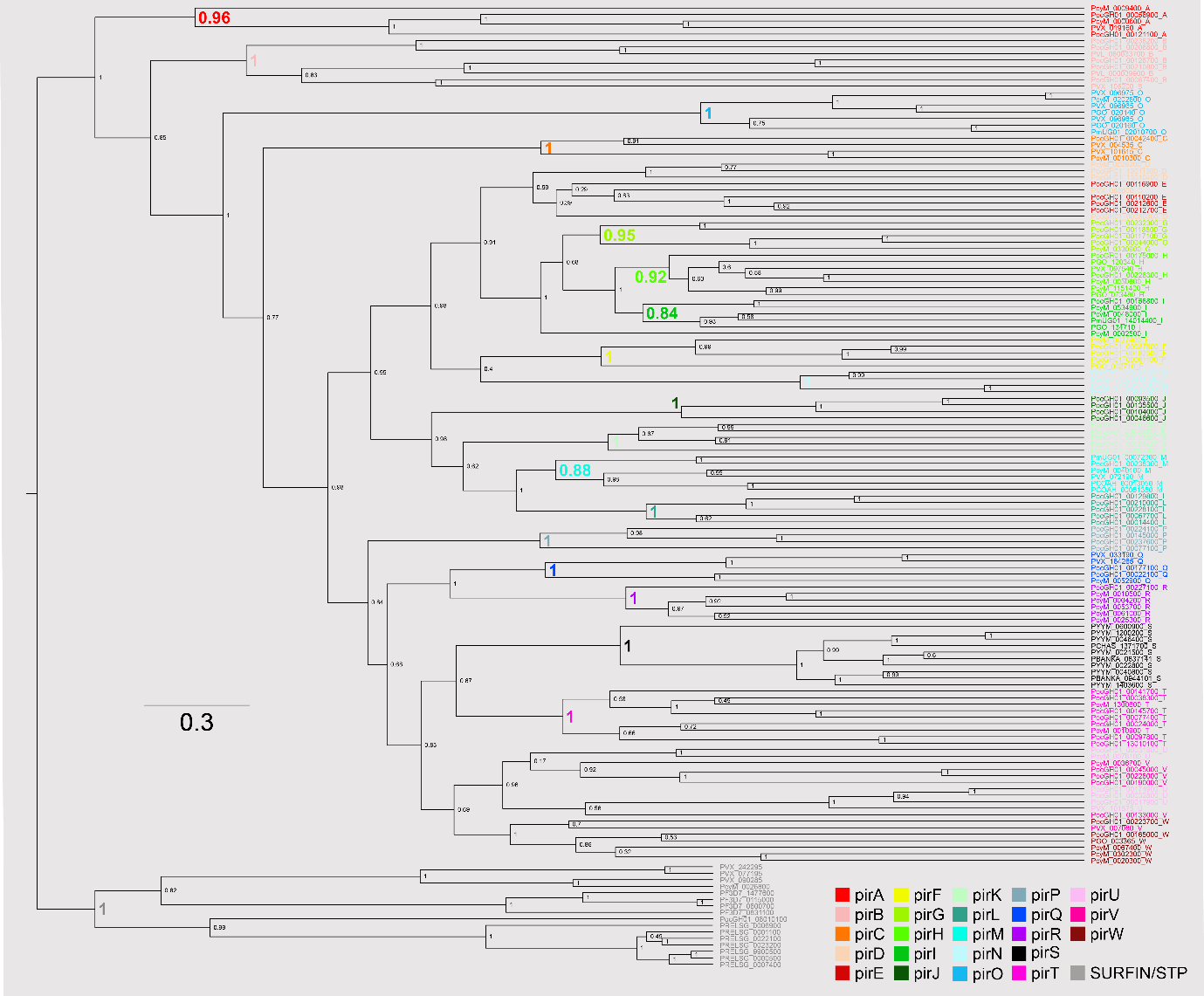

**Supplementary Figure 2. Bayesian phylogeny of *Plasmodium* interspersed repeat (*pir*) protein sequences.** The cladogram was generated using BEAST v2.7.7. from a 546-character alignment of 142 *pir* protein sequences taken from 15 *Plasmodium* or *Hepatocystis* species genomes, supplemented with 17 *surfin* protein sequences, which are designated as the outgroup. These sequences are a stratified subset of the larger dataset in Figure 1. Sequences are shaded by *pir* subfamily, as determined in Figure 1. Internode values show the Bayesian posterior probability; values subtending subfamily clades are shown in bold.

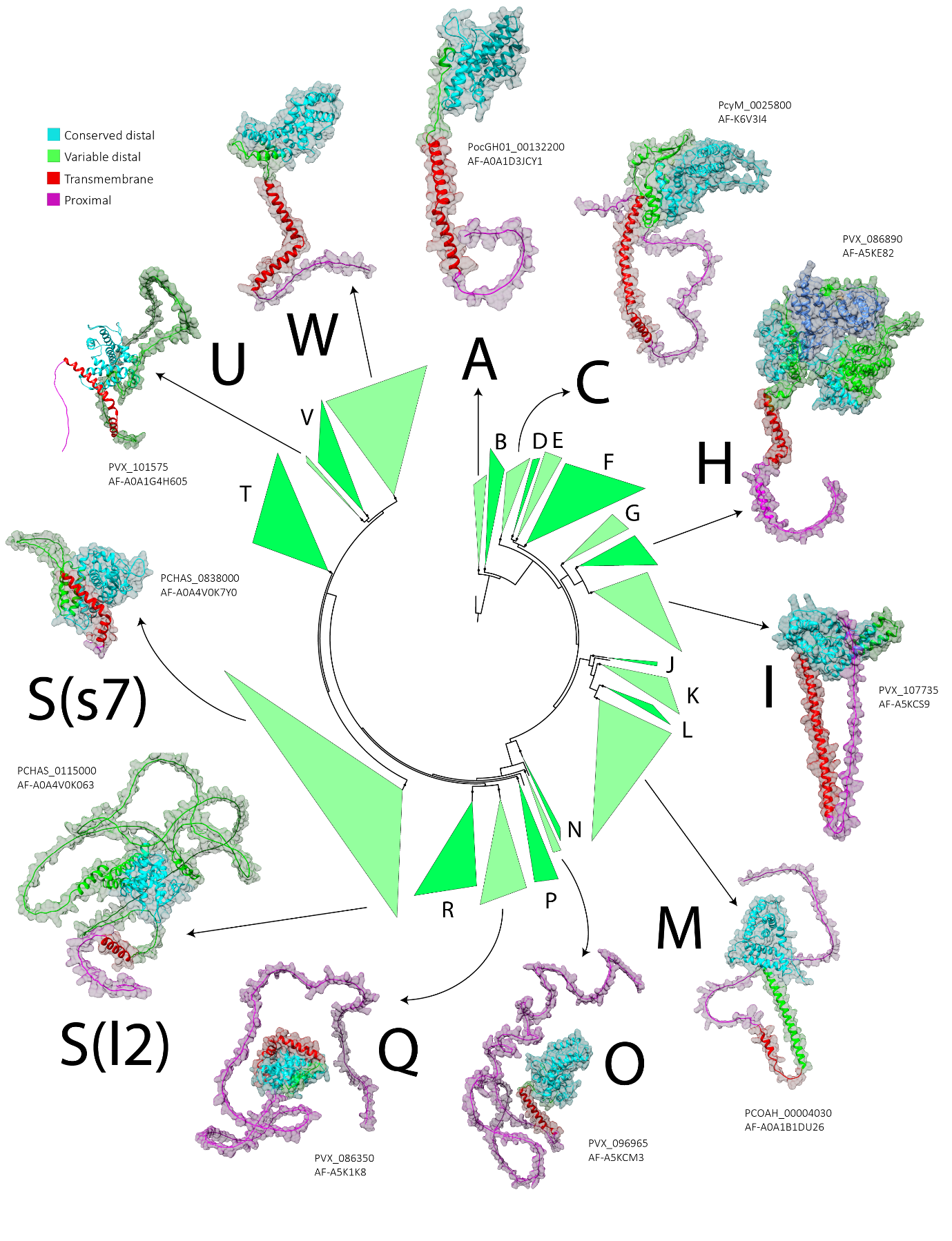

**Supplementary Figure 3. PIR protein structural variation**. Tertiary structures for single representatives of selected pir subfamilies predicted by Alphafold are shown in relation to a cartoon of the *pir* phylogeny. Each model is labelled with its PIR subfamily, specific PlasmoDB gene identifier and Alphafold identifier. Four regions of the PIR protein alignment, as defined in Fig. S1, are shaded for comparison. Some PIRH proteins contain multiple conserved cores, the additional core domain in the example here is shown in dark blue. Two examples are given for PIRS, one each for the short (‘s7’) and long (‘l2’) forms that are known among rodent PIR proteins.

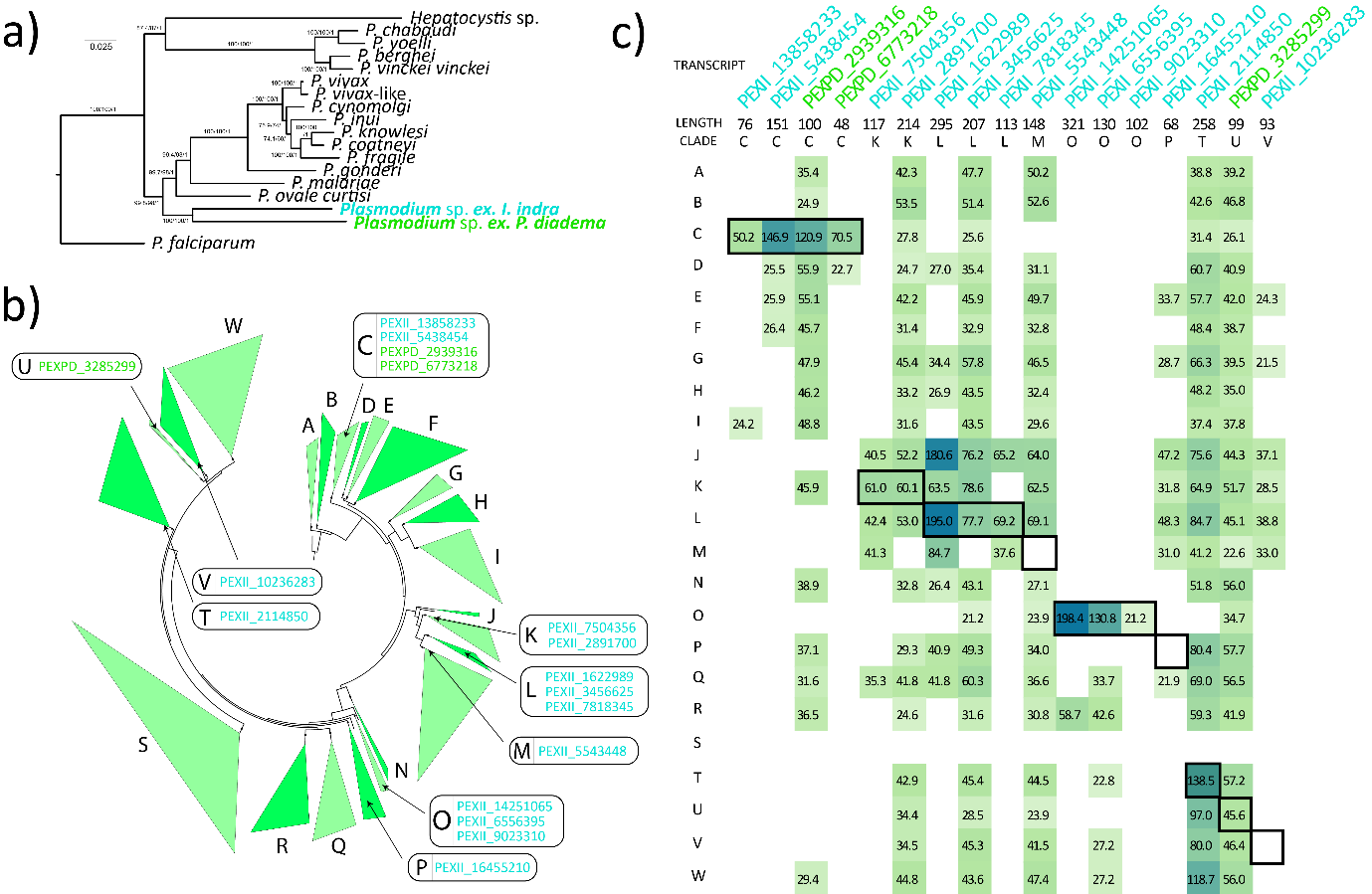

**Supplementary Figure 4. Lemur *Plasmodium* transcriptomes and *pir* repertoires.** Whole blood transcriptomes for two lemur species (*Indra indra*, *Propithecus diadema*) infected with malaria parasites were assembled from published RNA-seq data (see Methods). **A)** A phylogenomic tree showing the relationships of lemur malaria parasites and other species used in this study, generated from a concatenated alignment of 270 genes found in all species and estimated with both IQTREE and BEAST. Node labels show the maximum likelihood bootstrap value (n=1000) and posterior probability respectively. The tree is rooted with *P. falciparum*. **B)** A cartoon of the *pir* phylogeny (Fig. 1) showing the affinity of assembled transcripts from both lemur transcriptomes that are homologous to PIR according to BLASTx. **C)** Results of assessing structural similarity between predicted protein sequences of assembled transcripts and HMM models of each pir subfamily using HMMER v3.0. The HMM score for each transcript relative to each subfamily is reflected by the intensity of shading; highest scores are bounded.

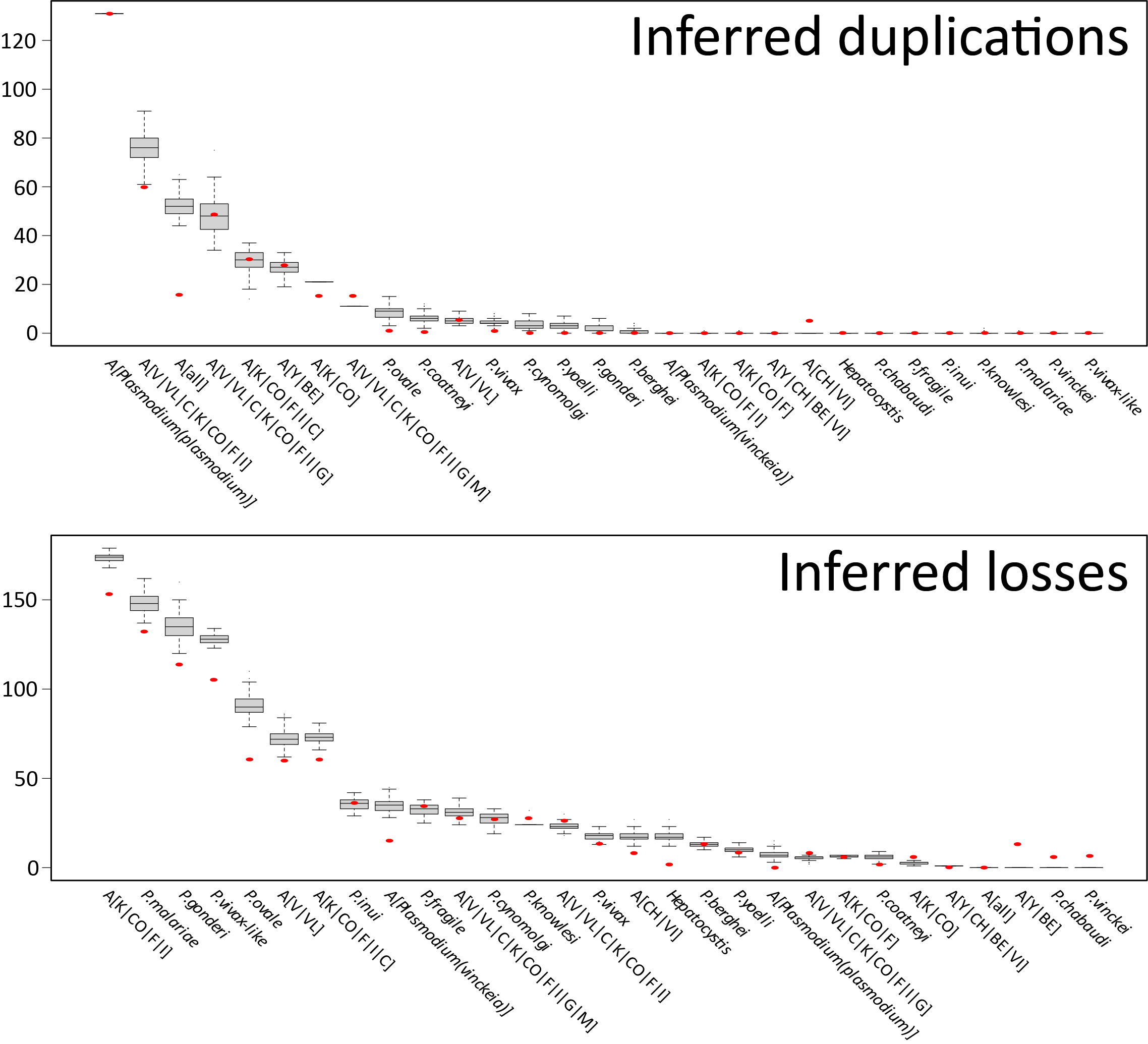

**Supplementary Figure 5. Effect of phylogenetic uncertainty on inferred gene duplication and loss frequencies.** Gene duplications and losses were inferred by reconciliation of gene and species phylogenies. Event numbers are dependent on tree topology, which is uncertain due to systematic error in phylogenetic estimation. To examine the effect of gene tree uncertainty on event numbers, 100 gene tree bootstraps were created and reconciled with the species tree. The range of values obtained are shown as grey box plots for each internal and terminal node. The value for the optimal gene tree is shown as a red dot. Internal nodes are described as ancestors (‘A[x]’) of their given daughter subgenera or species. Note that, while the bootstrap (suboptimal) gene trees necessarily introduce variation into the event numbers, the trend they produce is consistent with that obtained when the optimal gene tree topology is reconciled.

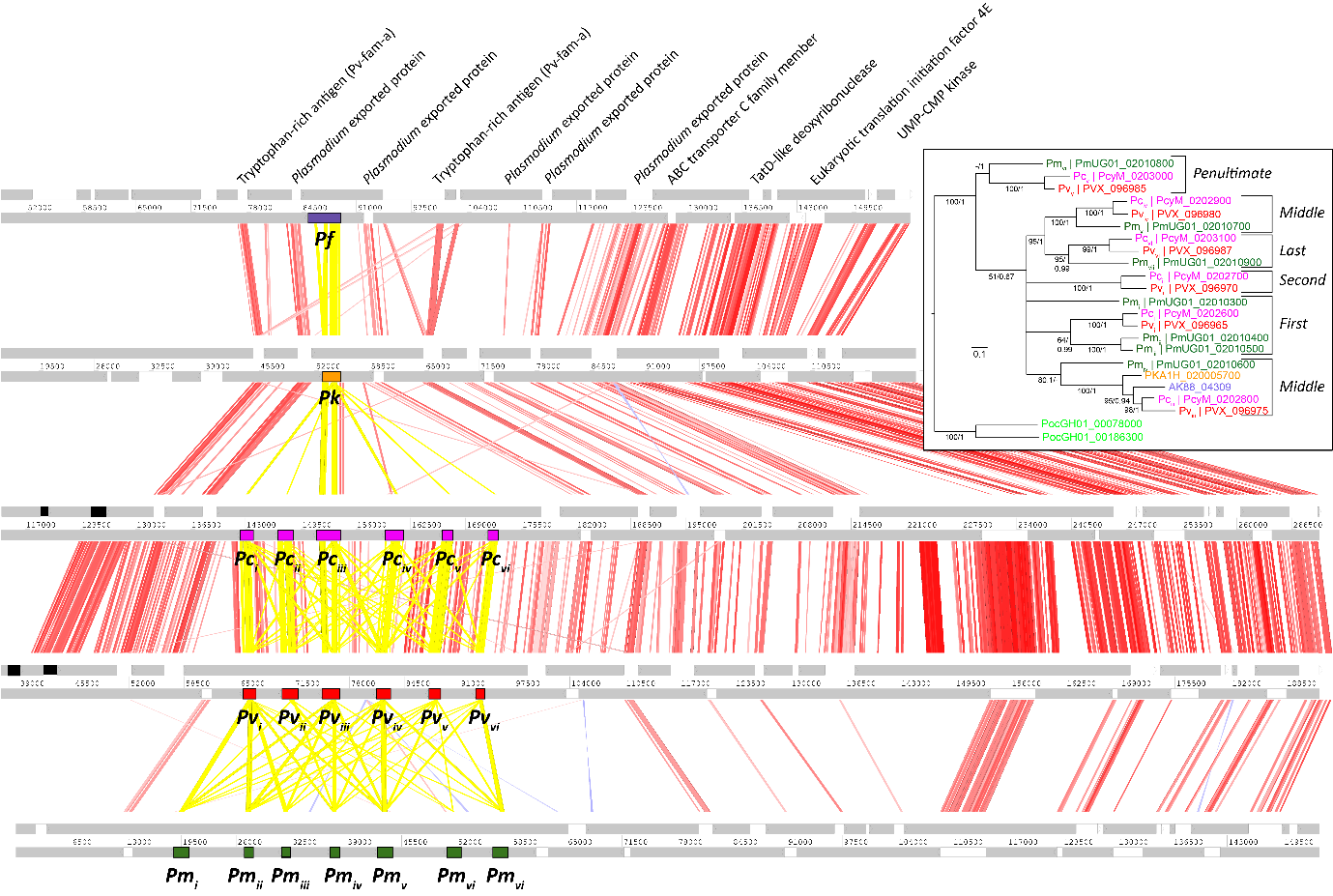

**Supplementary Figure 6. Conserved synteny of the *pirO1* locus across *Plasmodium* species.** *pirO* sequences from primate malaria genomes include orthologs that form a clade in the phylogeny. Those orthologs whose loci display conserved synteny at the core-subtelomere boundary on chromosome 2 are designated lineage *pirO1*. This locus is a tandem array of multiple between-species paralogs in several species. The structure of the locus is shown here for five species, (*P. fragile*, *P. knowlesi*, *P. cynomolgi*, *P. vivax* and *P. malariae*, from top to bottom); regions out of view to the left are subtelomeric while regions not shown to the right are core chromosome. Grey lines represent DNA strands and regions of sequence homology between genomes (assessed by tBLASTx) are shown as vertical red (non-*pir*) and yellow (*pir*) bars. Non-pir genes are marked by white rectangles and conserved non-*pir* flanking genes are labelled. *pir* genes are coloured and labelled by species; when arrayed, the position of *pir* genes within the array is indicated by subscript (*i* to *vii*), counting from left to right. Other *pir* genes (i.e. not *pirO1*) are shaded black. *Inset*: Maximum Likelihood phylogeny for the *pirO1* orthologs depicted based on an amino acid alignment rooted with *P. ovale* *pirO* sequences. Node values show bootstrap values and Bayesian posterior probabilities respectively. The tree shows that *pirO1* tandem copies retain orthology by position within the array across species. This indicates that both tandem duplication and structural differentiation of the copies occurred in the common ancestor of these primate malaria species and has been maintained (suggestive of purifying selection on discrete functions). In contrast, tandem paralogs have been lost from *P. fragile* and *P. knowlesi*, leaving just the ‘middle’ ortholog.

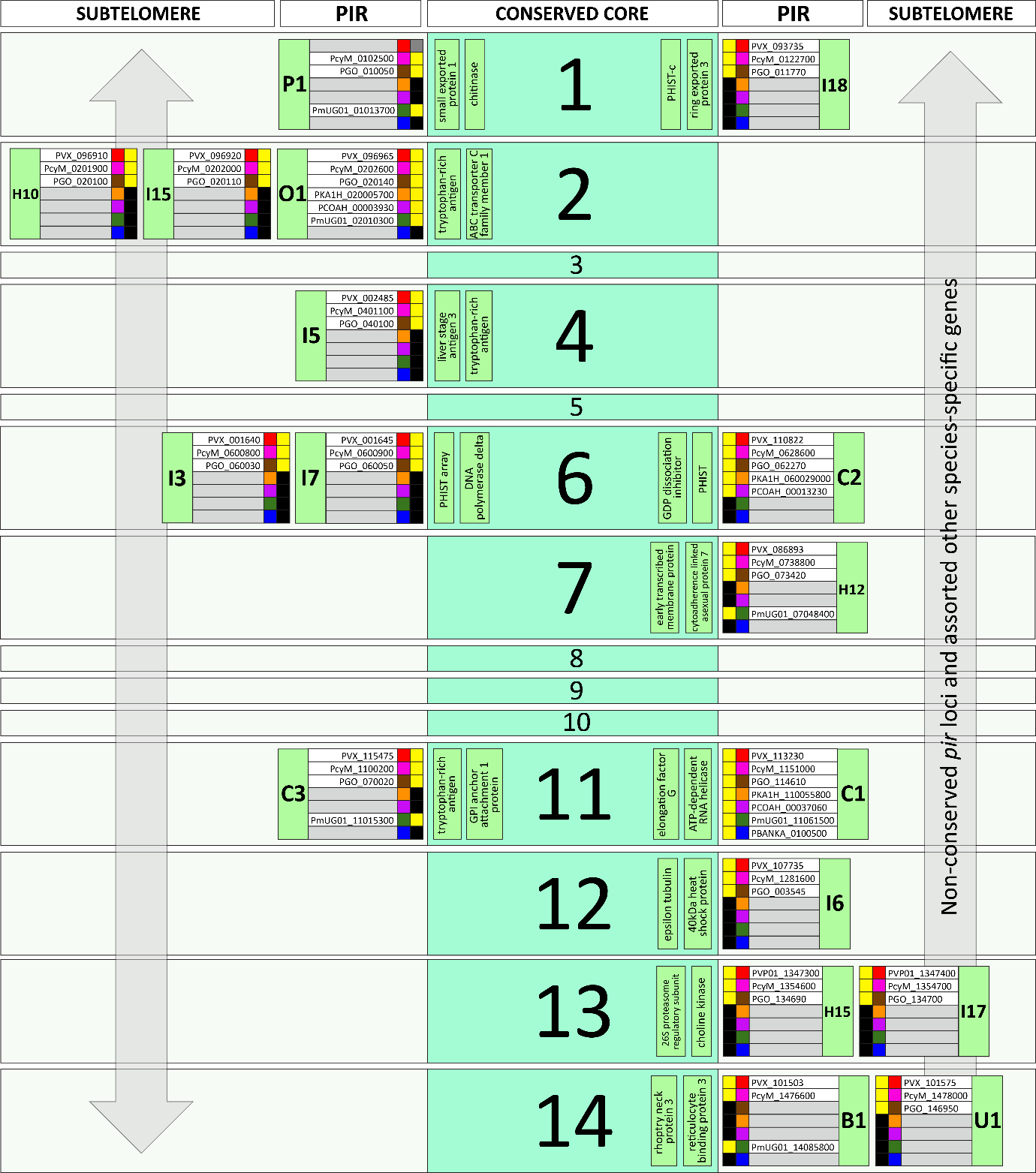

**Supplementary Figure 7. Chromosomal locations of conserved *pir* loci.** The cartoon presents the 14 chromosomes that are conserved across the *Plasmodium* species concerned here. A minority of *pir* loci are conserved across multiple species and these tend to occur at the boundary between the conserved chromosomal core (shaded green) and the non-conserved subtelomeric regions. These conserved *pir* genes are labelled after phylogenetic reconciliation (See Table S2). In each case, the presence (yellow) or absence (black) of the locus from seven genomes is noted (*P. vivax*, red; *P. cynomolgi*, pink; *P. gonderi*, brown; *P. knowlesi* (orange), *P. coatneyi* (purple), *P. malariae* (green), *P. berghei* (blue)). The subtelomeric regions are sufficiently contiguated in these seven genomes to make an analysis; note that these loci will be present in additional species (e.g. *P. ovale*) whose genomes do not currently represent their subtelomeres. When present, the PlasmoDB identifier for the orthologous gene is given. Also, selected, conserved genes that help to define the position of the *pir* locus as shown within the core region.

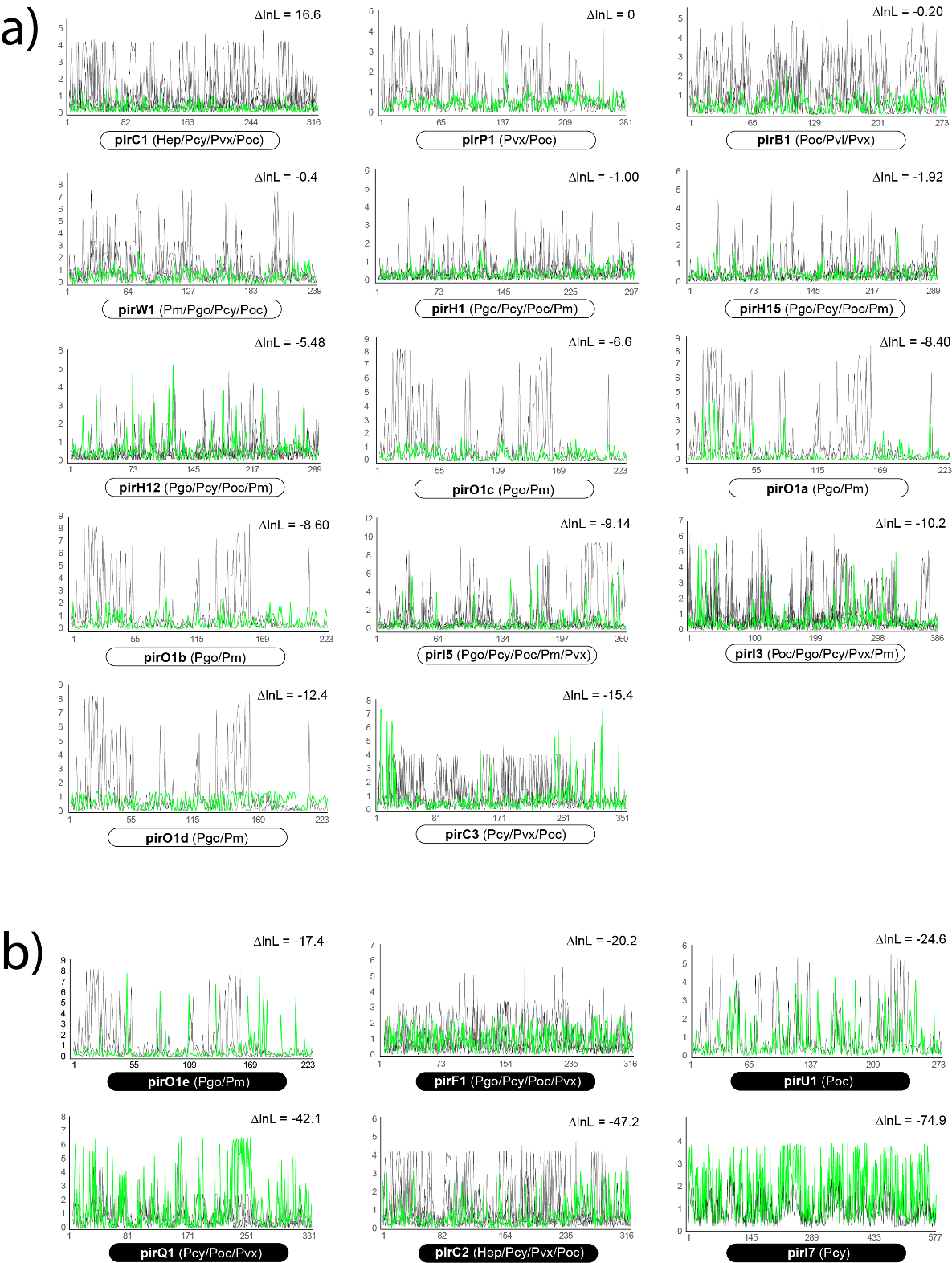

**Supplementary Figure 8. *D_n_/D_s_* (ω) ratio values compared for orthologous and paralogous codon sequences across multiple *pir* subfamilies.** *D_n_/D_s_*, the ratio of synonymous to non-synonymous amino acid substitutions, (ω) was estimated using codeML for codon alignments of orthologous *pir* sequences. Orthology was inferred by reconciliation analysis of the *pir* phylogeny and confirmed with comparison of genomic context (see methods). Each graph plots ω against position within the codon alignment for the ortholog (green) and for one or more paralogs (black). The species from which paralogs were sampled are noted below the x-axis. a) *pir* orthologs that have experienced significant negative (purifying) selection in contrast to their related paralogs; codon models with and without negative selection show a significant difference in -lnL values (typically > -16; see text). b) *pir* orthologs that have not experienced negative (purifying) selection and show no significant difference with their related paralogs.

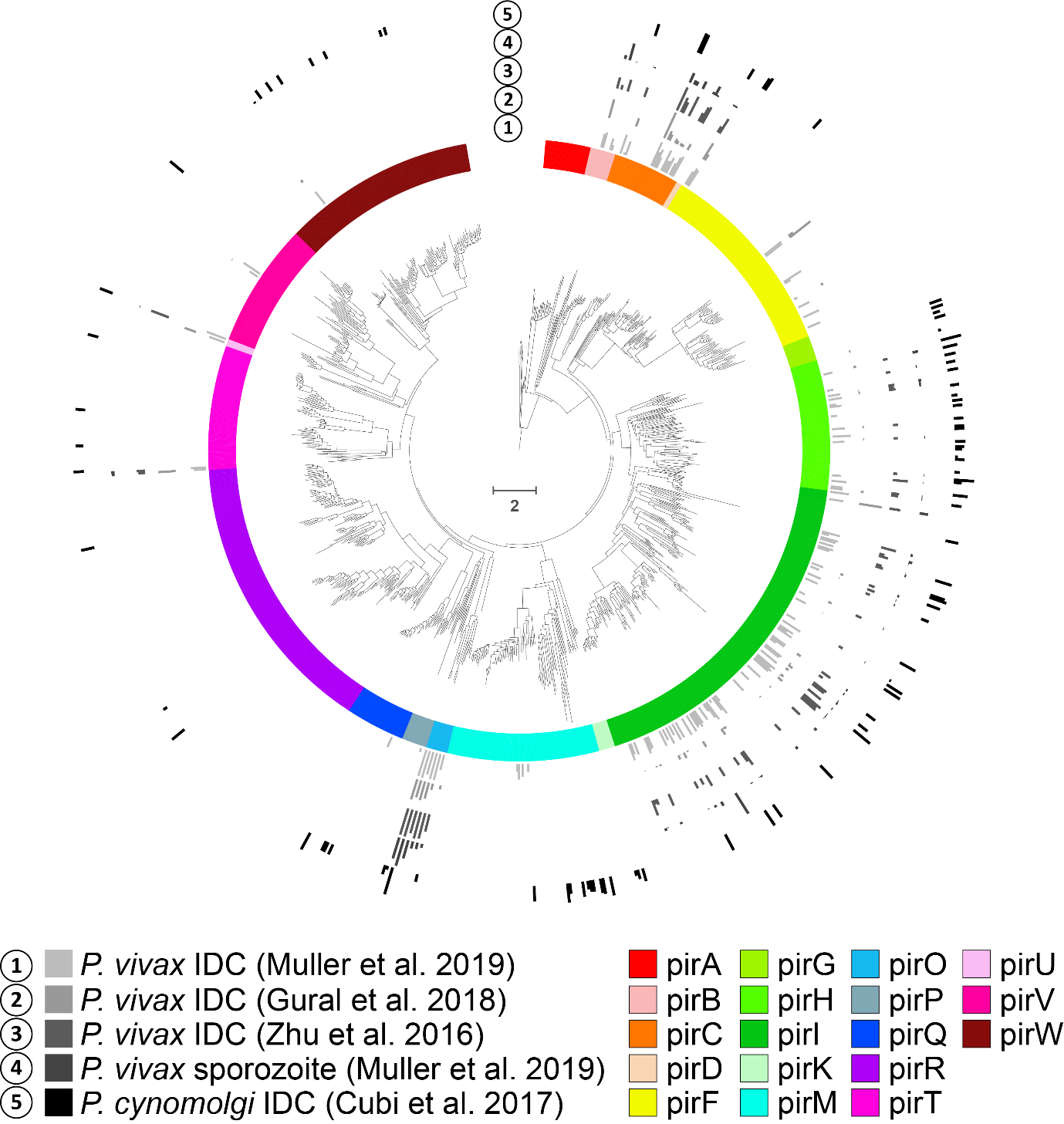

**Supplementary Figure 9. Transcript abundance for expressed *P. vivax* and *P. cynomolgi* *pir* genes mapped to phylogeny.** The central phylogram shows the relationships of *P. vivax* and *P. cynomolgi* *pir* genes extracted from the *pir* phylogeny (Figure 1); *pir* subfamilies are shaded by colour. Five tracks are arranged around the outside, each displaying transcript abundance values from a published parasite transcriptome: 1) intraerythrocytic bloodstream stages of *P. vivax* (Muller et al. 2019); 2) intraerythrocytic bloodstream stages of *P. vivax* (Gural et al. 2018); 3) intraerythrocytic bloodstream stages of *P. vivax* (Zhu et al. 2016); 4) *P. vivax* sporozoites (Muller et al. 2019); and 5) intraerythrocytic bloodstream stages of *P. cynomolgi* (Cubi et al. 2017). The height of the bar aligned to each gene is proportional to transcript abundance.

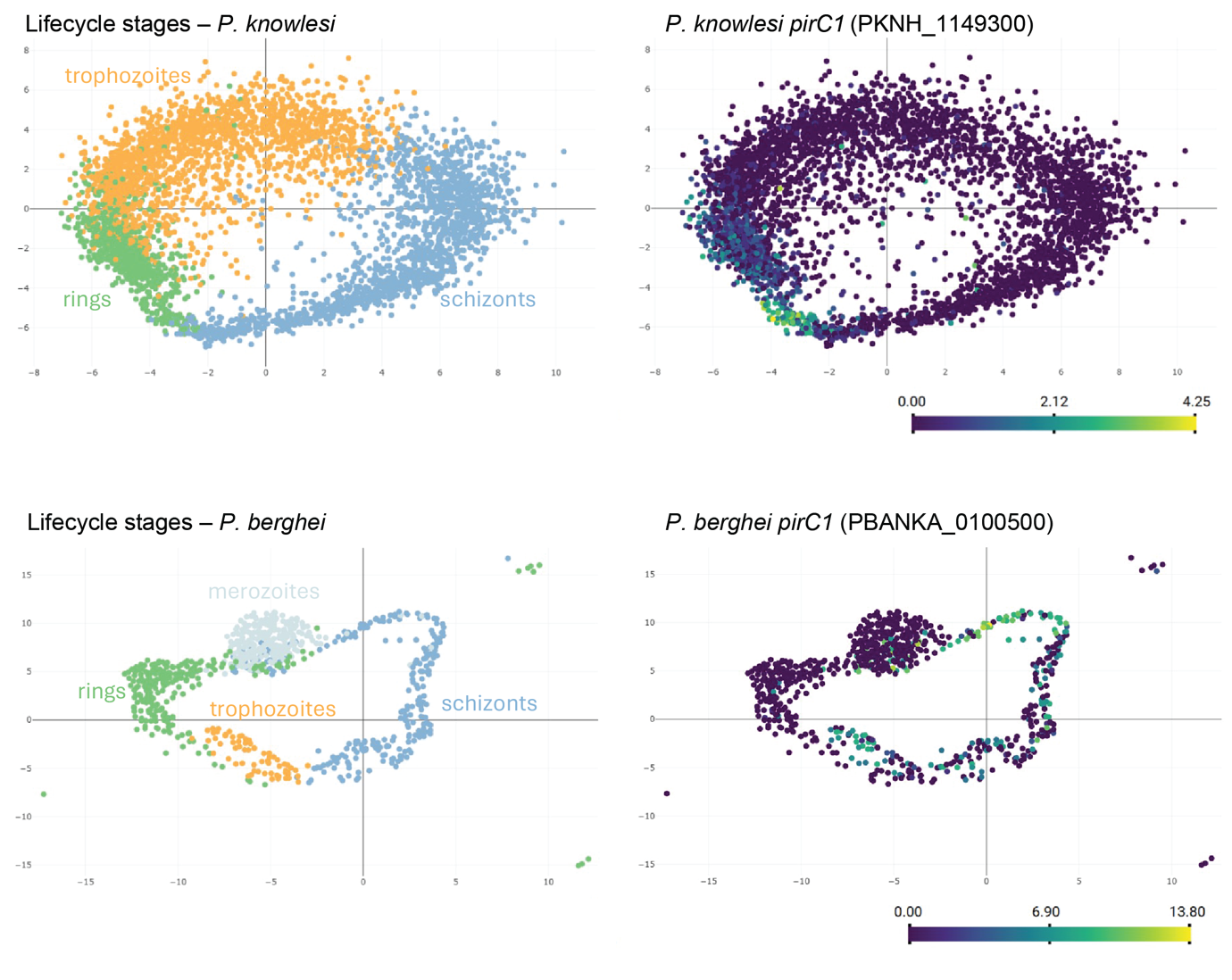
**Supplementary Figure 10. Expression profiles of *pirC1* in *Plasmodium knowlesi* (top) and *Plasmodium berghei* (bottom).** Panels on left-hand side show life cycles stages, panels on the right show the expression of the *pirC1* genes. Data obtained from the Malaria Cell Atlas (Howick *et al*, 2019) .

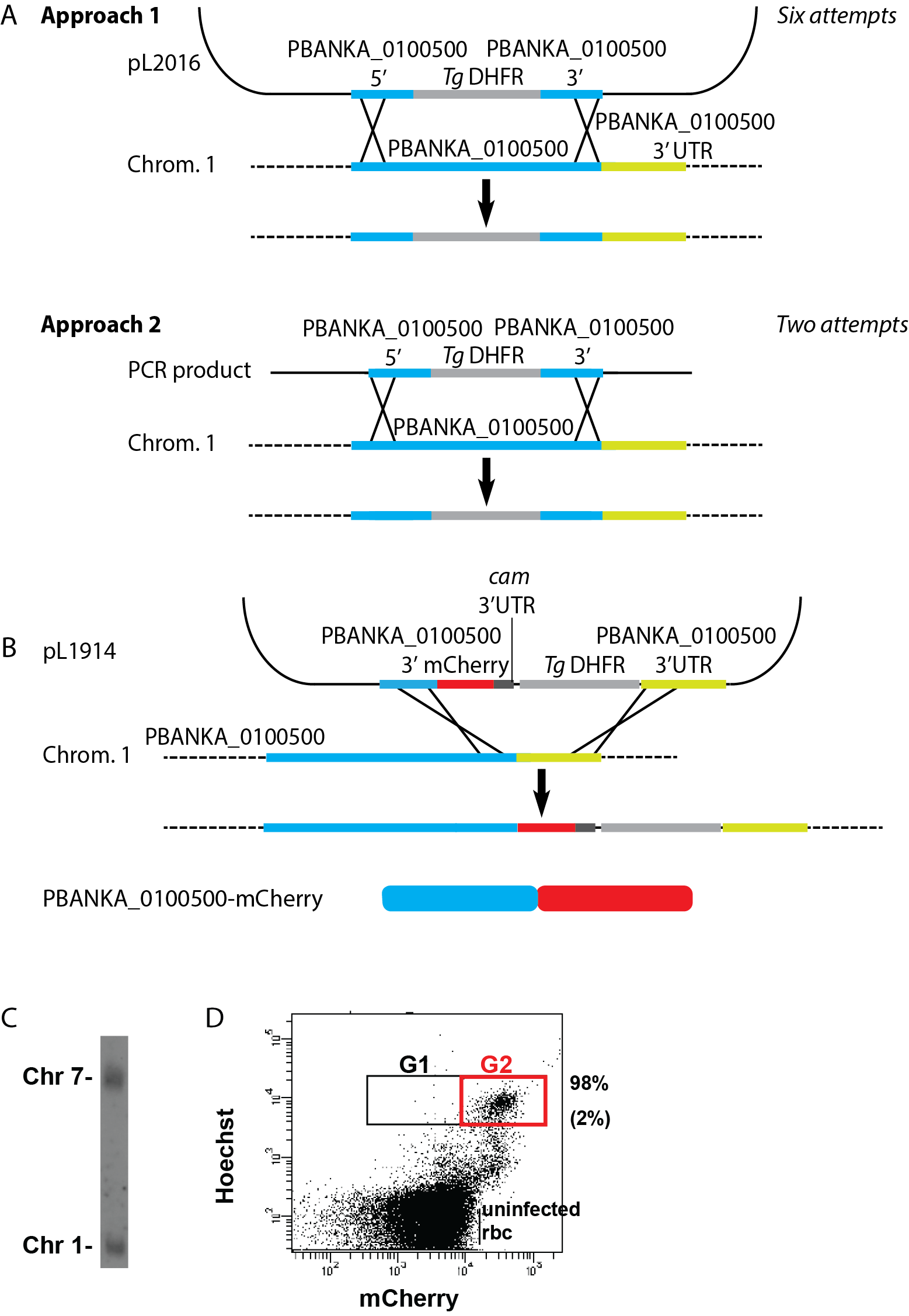

**Supplementary Figure 11. Genetic modification of *Plasmodium berghei*. A.** Schematic representation of the introduction of the selectable marker (SM) cassette into the *PBANKA_-0100500* gene locus of the wild type ANKA *P. berghei* parasites. Upper panel, construct pL2016 contains the SM (*tgdhfr/ts* selectable marker cassette) flanked by *bir* target regions and integrates into the *bir-0100500* locus by double cross-over homologous recombination. Plasmid is linearized at the Asp718/XbaI sites. Lower panel, PCR-construct, pL2029, contains the SM (*hdhfr* selectable marker cassette) flanked by *bir-0100500* target regions and integrates into the bir-0100500 locus by double cross-over homologous recombination. **B.** The construct (pL1940) used for generation of transgenic line BIR-0100500::mCherry that expresses *bir-0100500* C-terminally tagged with mCherry. The construct contains the *tgdhfr/ts* selectable marker cassette (SM) and a target region of *bir-0100500* for integration by single crossover homologous recombination after linearization with the BsmI site. Integration of the construct into the *bir-0100500* locus results in a C-terminal mCherry-tagged copy of *bir-0100500*. B. Integration of **C.** Genotype analysis of BIR-0100500::mCherry confirming correct integration of pL1940. Southern analysis of separated chromosomes shows integration of the tagging construct in chromosome 1, using the 3′utr *dhfr* probe which hybridizes in the SM cassette of pL1940 and the endogenous dhfr/ts locus (chromosome 7). **D**. Flow cytometry analysis of BIR-0100500::mCherry schizonts. The O/N cultured infected red blood cells were selected based on Hoechst and mCherry fluorescence. An average percentage of 98% (sd 2%; n = 3; Gate 2) of the total number of schizonts (Gate 1) were mCherry positive.

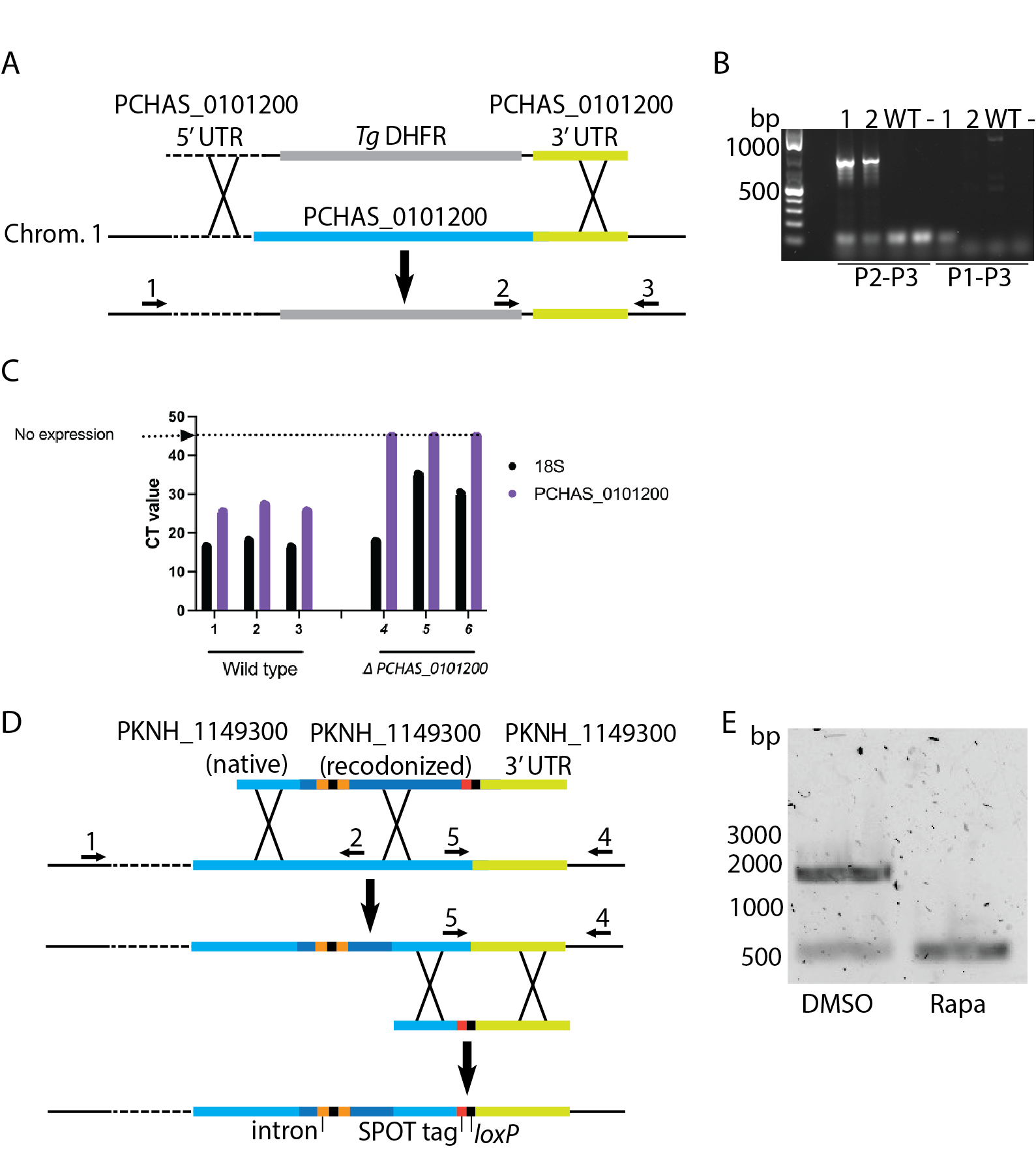

**Supplementary Figure 12. Genetic modification strategy and genotyping of *Plasmodium chabaudi* and *Plasmodium knowlesi* parasites**. A) *Plasmodium chabaudi* AS parasites lacking PCHAS_0101200, (Δ*PCHAS_0101200*) were generated by targeting the PCHAS_0101200 locus with the *P. chabaudi* AS *ΔPCHAS_0101200* construct, in which the *P. chabaudi smac* region in the *Pc*AS Δ*smac* construct described previously (Cunningham *et al*, 2017) was replaced with the *P. chabaudi* PCHAS_0101200 targeting region (PCHAS_01_v3 50360-49650 and PCHAS_01_v3 52515-51810), followed by transfection of with the Kpn1-SacII digested plasmid. B) Integration into the PCHAS_0101200 locus was verified by PCR using primers P2 (within the plasmid) and P3 (downstream of the integration site). PCR products from the ΔPCHAS_0101200 parasites (1-2), wild-type *P. chabaudi* AS parasites (WT) and H_2_O control (‑), amplified with primer pairs P2-P3 and P1-P3, as indicated. The integration product is only present in the ΔPCHAS_0101200 line (1 and 2) and the wildtype product in the control line only (WT). C) qRT-PCR amplification of the ΔPCHAS_0101200 and wild-type *Pc*AS parasites showed the absence of ΔPCHAS_0101200 transcription in the ΔPCHAS_0101200 line after 45 amplification cycles (CT value=45). Control parasites exhibited high expression of PCHAS_0101200, with CT values of 25-27. All samples expressed the 18S housekeeping (PCHAS_0937240) gene (CT values of 17-34 and 16-18). D. Modification of *Plasmodium knowlesi* parasites. The native PKNH_01149300 locus was targeted with a PCR product to introduce two *loxP* sites within and following the gene. However, initially only parasites with a single *loxP* site were obtained owing to recombination between the wildtype and recodonized sequence. The second *loxP* site was introduced using Cas9-mediated integration of pBLD700. This also introduced sequence encoding the SPOT tag into the gene. E) Removal of the gene was verified by PCR with primer pair 1-5 using genomic DNA from parasites treated with DMSO (the vehicle control) or Rapamycin. Note that even in DMSO-treated parasites, a low level of excision was detected.

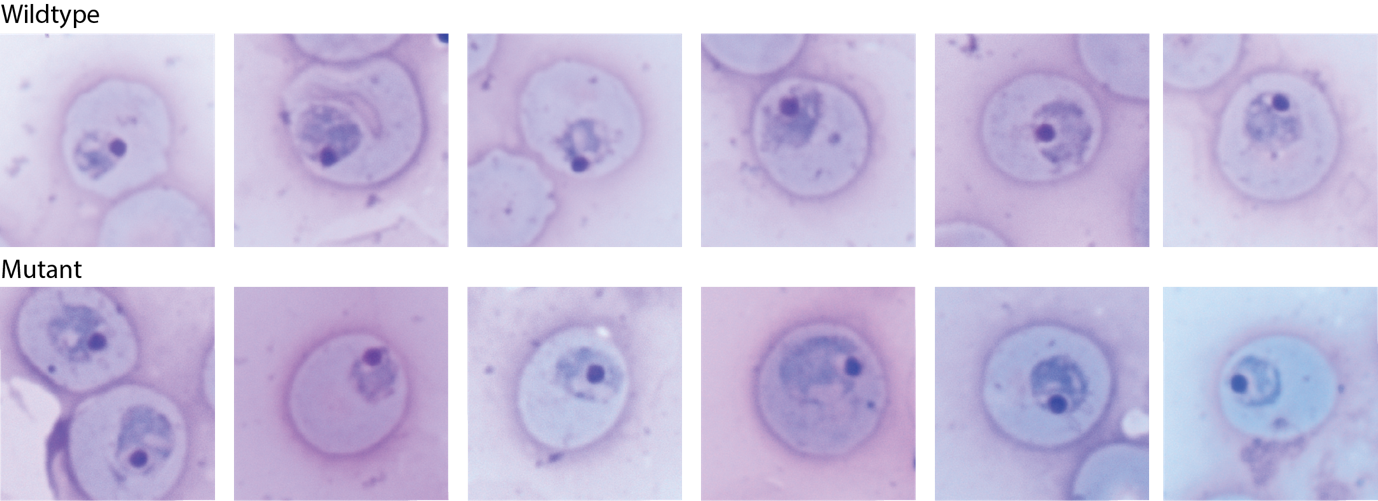

**Supplementary Figure 13. Additional examples of *Plasmodium chabaudi* wildtype parasites and parasites lacking PIRC1.** These images were obtained from the same experiment as the images shown in Figure 4B.

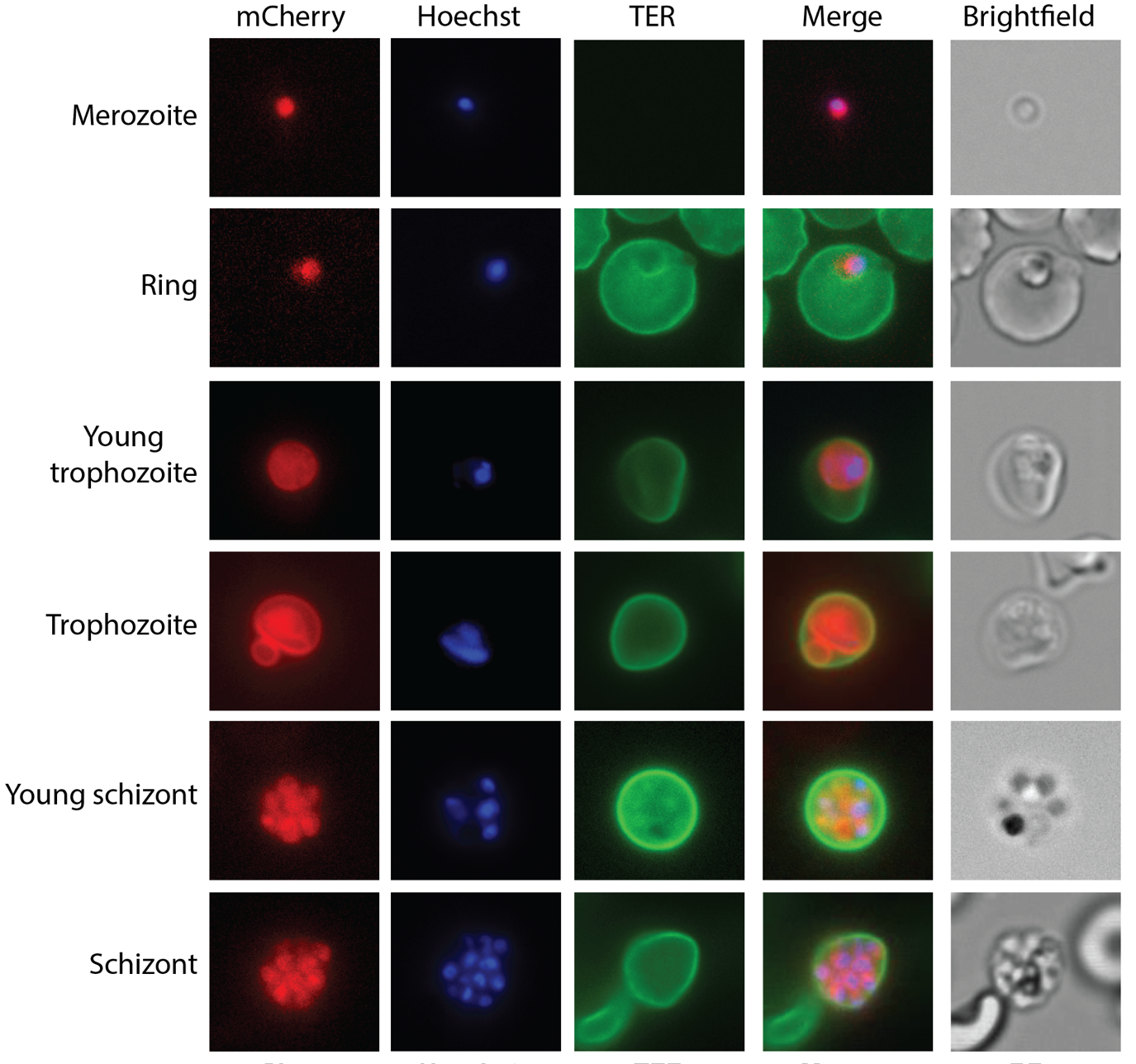

**Supplementary Figure 14. Localization of *Plasmodium berghei* *PirC1*-mCherry.** These images were obtained in the same experiment as the images shown in Fig. 4C.

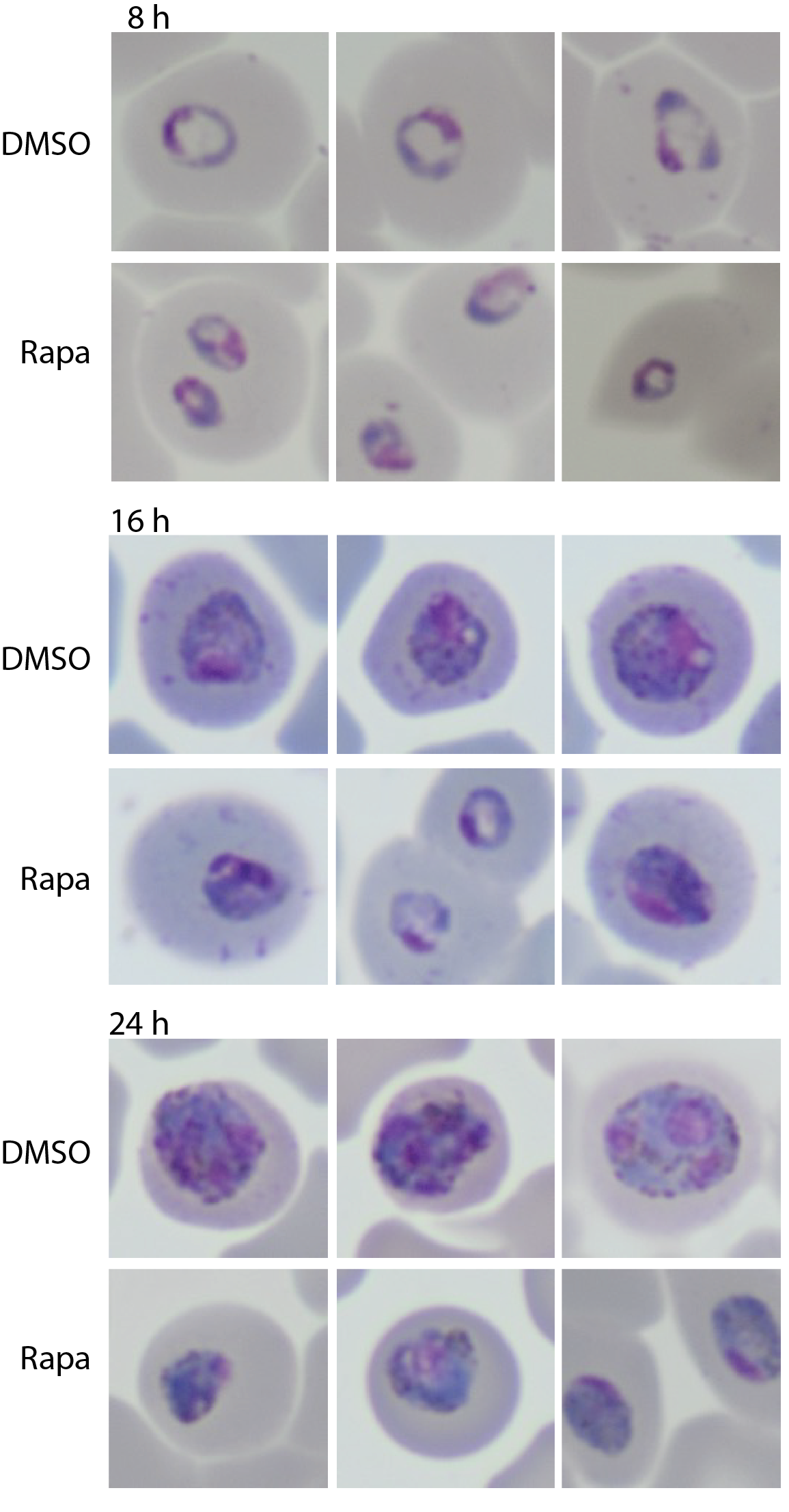

**Supplementary Figure 15. Additional examples of wildtype parasites and *Plasmodium knowlesi* parasites lacking *pirC1* at different times after invasion.** These images were obtained from the same experiment as the images shown in Fig. 5B.

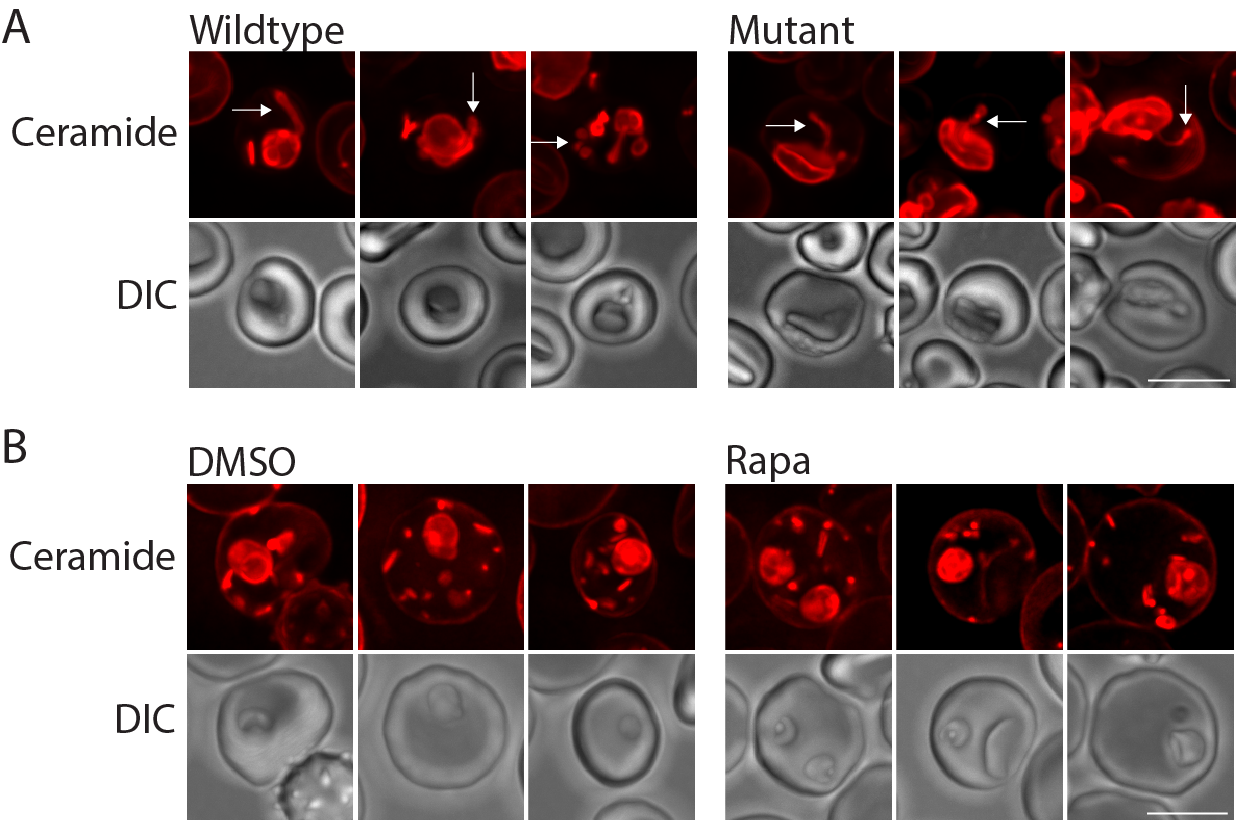

**Supplementary Figure 16. The exomembrane system in erythrocytes infected wildtype *Plasmodium knowlesi*** **parasites or parasites lacking *pirC1*.** (A) Wildype *P. knowlesi* parasites (DMSO) and parasite lacking *pirC1* (Rapa) (B). The exomembrane system was visualized by staining the cells with C5-Bodipy-ceramide. The cells were imaged live. Arrows in top panels indicate exomembrane system. Note that there is no obvious difference in the exomembrane system of the wildtype parasites and the parasites lacking *pirC1*. The scale bar represents 5 μm.

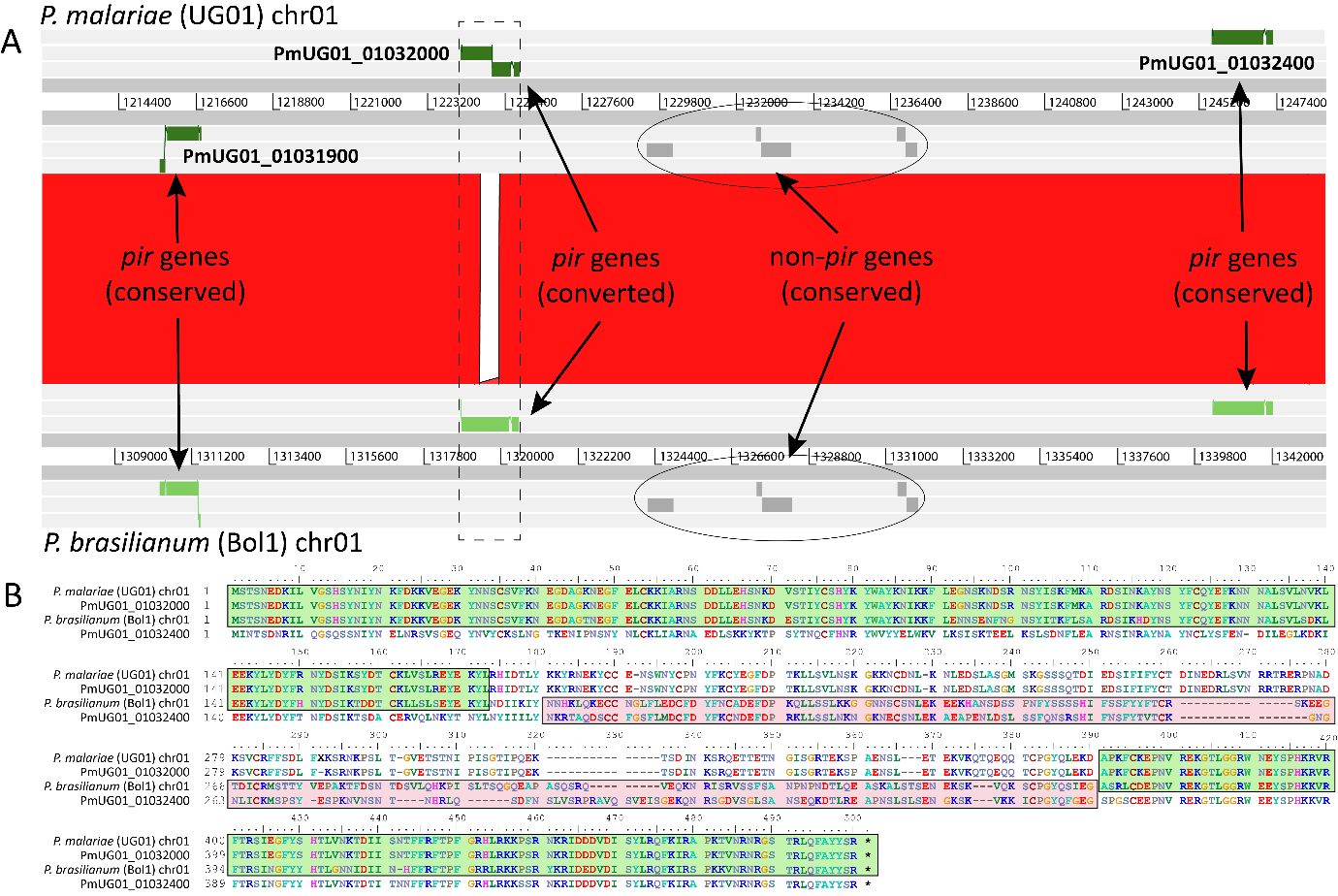

**Supplementary Figure 17. Alignment of a homologous subtelomeric region of chromosome 1 from *P malariae* (UG01) and *P. brasilianum* (Bolivia1).** A. Genomic comparison of three subtelomeric *pir* loci. Grey lines represent DNA strands and regions of sequence homology between genomes (assessed by BLASTn) are shown as vertical red bars (97-99% nucleotide identity). *Pir* genes are shaded green; non-*pir* genes are marked by grey rectangles. The only region that does not show within-species levels of sequence conservation is the middle portion of the second *pir* locus (PmUG01_01032000). The closest match in *P. malariae* to the *P. brasilianum* sequence homoeologous to PmUG01_01032000 is the third *pir* gene shown here (PmUG01_01032400). B. Sequence alignment of the regions within the dashed lines with the PmUG01_01032000 and PmUG01_01032400 transcripts. This shows how the *P. brasilianum* sequence is conserved with *P. malariae* at the 5’ and 3’ ends of the gene (green shading), but departs in the central portion, where it is more alike to PmUG01_01032400 (red shading). Since the agreement between *P. brasilianum* and PmUG01_01032400 is not very close, we may suggest that another gene was the donor for gene conversion, but that gene is not present in the *P. malariae* UG01 assembly. Gene conversion events of this kind explain how pir sequences can be duplicated and deleted rapidly within individual genomes.

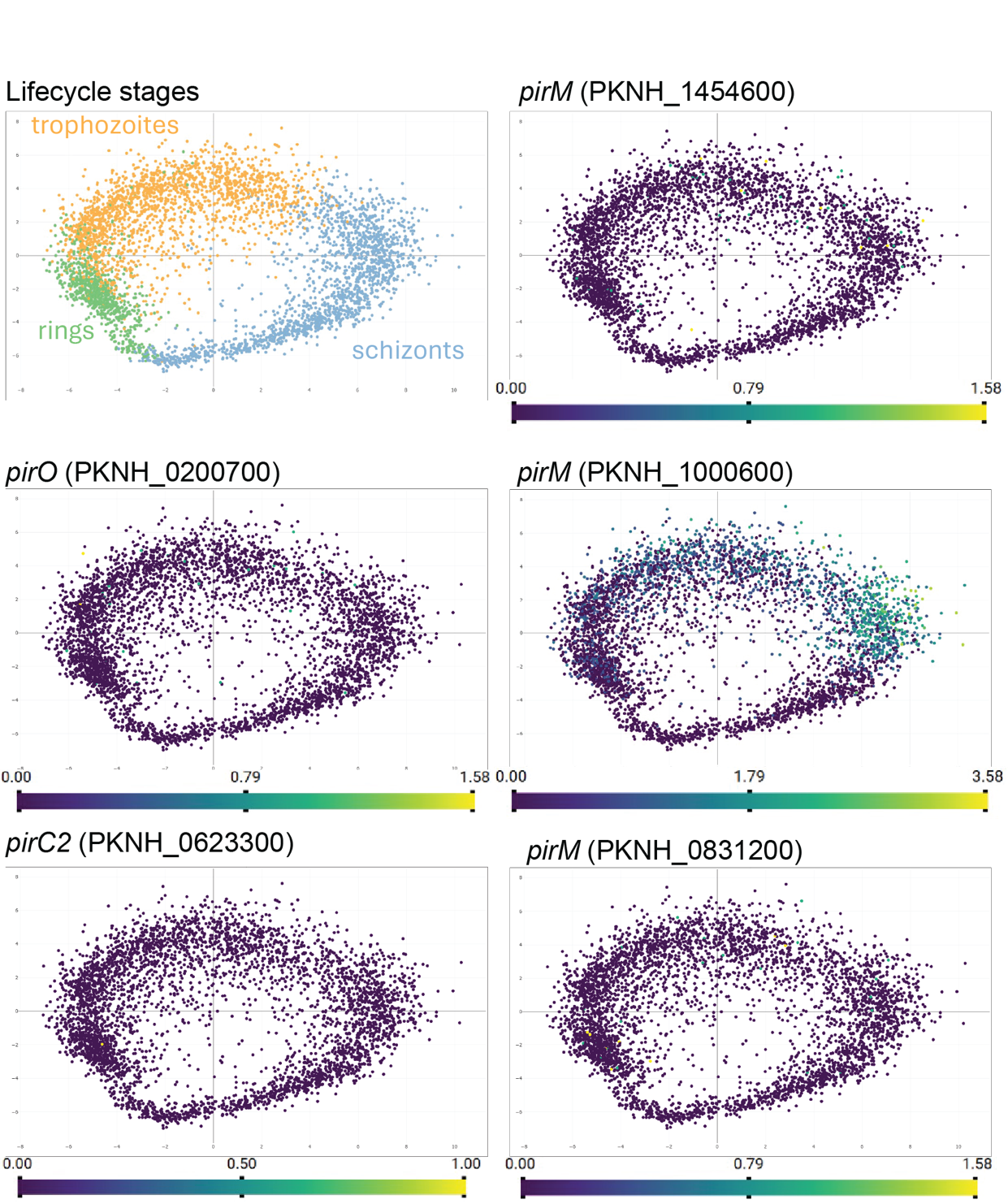

**Supplementary Figure 18. Expression of selected *Plasmodium knowlesi* *pir* genes.** Note the low expression of the *pirO* and *pirC2* genes, whereas the *pirM* gene PKNH_1000600 is expressed at a high level in all or nearly all cells in the population.
