## Supplementary results for "Essential function reflected in the phylodynamics of a multigene family – the *pir* genes of malaria parasites"

**Multiple sequence alignment and PIR structure**

The multiple alignment we created to estimate *pir* phylogeny contains 4232 amino acid sequences and 839 aligned positions. The alignment recapitulates what is known about *pir* structure (del Portillo *et al*, 2001; Janssen *et al*, 2004; Frech & Chen, 2013), as illustrated in Figure 3a. There is a core of seven alpha helices that contains multiple, conserved cysteine residues that make it easily alignable. This corresponds to exons 1 and 2 of *pir* genes and approximately the first 500 positions of the alignment. As a transmembrane protein, this domain is predicted to be furthest from the parasite nucleus when expressed *in situ*; hence, we refer to this here as the ‘distal conserved’ domain. The transmembrane domain is also conserved and, together with its immediate flanking regions, approximates to positions 510-710 of the alignment. In PIR proteins, the sequence between the conserved alpha helices and the transmembrane domain is highly variable and can be quite short (e.g. *pirO*) or extremely long (e.g. *pirS*). Hence, we refer to this as the ‘distal variable’ domain. Typically, it cannot be aligned except among the most closely related genes and so was largely removed from the alignment; we shortened it to between 20 and 40 amino acids prior to the transmembrane helix to provide clade-specific characters. The domain following the transmembrane helix corresponds to positions 710-839 of the alignment. This domain is predicted to be intracellular and so closest to the parasite nucleus; hence, we refer to this part as the ‘proximal’ domain. Like the distal-variable domain, this can be quite short (e.g. *pirU*) or very long (e.g. *pirQ*) and is clade-specific everywhere. Since it is not alignable across all *pir*, (except perhaps for the extreme C-terminus) it was largely removed. We retained the first 150 amino acids adjacent to the transmembrane helix to provide useful clade-specific characters within subfamilies. Note that the distal-conserved and transmembrane domains are shared with SURFINs.

**The *surfin* genes are the natural outgroup to *pir* and its likely progenitor**

The structural similarity between *pir* and *surfin* was described by Frech and Chen (Frech & Chen, 2013). Both proteins are predicted to have a single transmembrane domain and their extracellular domains (corresponding to the second exon in *pir*) are clearly and easily alignable due to the conserved pattern of cysteine residues that they share. *Surfin* and *pir* may be distinguished by the much larger intracellular component of *surfins* and distinctive first exon of *pir*. When searching genome sequences for *pir* with BLAST or HMMER, it is usual to also identify *surfin* sequences after the least related *pir*. There are no *pir* sequences in *P(Laverania)* genomes or *P(Haemamoeba*) such as *P. relictum, P. gallinaceum.* When searching the *P. falciparum* genome using BLASTp with a PIR query, no PIR are found but SURFIN is the closest match. Beginning with this *P. falciparum surfin* sequence (i.e. PF3D7_0113100), we identified all related sequences across all available *Plasmodium* genomes. This identified a *surfin* clade including sequences from *P. falciparum, P. gaboni, P. relictum, P. gallinaceum*, *P. ovale, P. vivax, P. cynomolgi* and *P. malariae*, which is sister to another clade of ‘STP’ sequences from *P. ovale, P. vivax* and *P. malariae*. Combined, these two clades comprise 56 sequences that represent the most closely related sequences to *pir* that lack the conserved structural dimensions of *pir*. In both maximum likelihood and Bayesian analyses, these sequences robustly form a clade to the exclusion of all canonical *pir*. Since *surfin* genes are found across *Plasmodium*, they have a wider distribution than *pir* genes, (which are restricted to *P*(*Plasmodium*) and *P*(*vinckeia*) subgenera). All else being equal, this means that *surfin* are older than *pir* and, at present, the likely origin of *pir* genes is from a *surfin*-like sequence in the common ancestor of *P*(*Plasmodium*) and *P*(*vinckeia*).

**Tandem duplication of the ‘core’ domain in some *pir* genes**

Sequence alignment revealed that it is quite common for *pir* genes in certain sub-families (primarily *pirH*, *pirI* and *pirM* but also *pirQ*) to contain a tandem duplication (or triplication) of the distal-conserved domain (i.e. the first seven alpha helices). We refer to this phenomenon as ‘multiple cores’. This might occur if multiple *pir* genes arranged in tandem had been combined in error. Upon inspection, most of these affected sequences were deemed accurate because the multiple cores did not possess multiple transmembrane domains, which they would if gene models had been falsely spliced together. 94 sequences have a triplicated core, while 170 have a duplicated core; these amount to 6.26% of all *pir* aligned (see Data File 3). Ten sequences (8 x *P. coatneyi* *pir*, 1 x *P. knowlesi* and 1 x *P. yoelli*) were found to be errors and removed, including one gene model (PCOAH_00003180) that falsely combined seven or eight separate *pirs*. In the alignment, only the core duplicate closest to the transmembrane domain was retained for analysis.

**Bayesian phylogenetic analysis supports the designated *pir* subfamilies**

To further evaluate the accuracy of the maximum likelihood phylogeny, a Bayesian phylogenetic analysis of 142 sequences drawn from all sub-families was conducted using BEAST. The tree sample was examined in Tracer, which showed that the MCMC was well mixed and converged on a robust topology. Thus, mean treelength was 168.4 and had an ESS of 3904, while mean tree Likelihood was -102122.4 and had an ESS of 2149. Estimates of both parameters displayed a normal distribution. Figure S1 shows that clades designated as sub-families in the maximum likelihood topology are usually reproduced with a high posterior probability (pp) approaching one. *PirI* (pp=0.84) and *pirM* (pp=0.88) are monophyletic but with lower support. Exceptions are *pirD*/*E* and *pirW/U/V*; in these two situations, the sub-families are paraphyletic, typically with their sister clades as defined in Figure 1. Hence, there is minor uncertainty about the monophyly of *pirD*, *pirE*, *pirU* and *pirV*. Sequences that branch basally within these families have uncertain affinity. The Bayesian topology reproduces the robust reciprocal monophyly of *pir* and *surfin*/STP sequences. With respect to supra-familial relationships, some larger clades are well supported (e.g. *pirD-N* and *pirS-W*), and *pirA* remains the most basal-branching lineage, but these deep nodes that relate sub-families are not always robust, so the basal topology can be expected to change with the addition of more sequences or the application of more precise models.

**Transcriptomes from lemur malaria parasites demonstrate the utility of a universal *pir* systematics**

A practical outcome of a comprehensive comparative analysis is a universal and exhaustive systematics for *pir* genes, one that might be applied to any species, past, present or future, and which allows us to speak about the same *pir* gene in different species. The subfamilies established here should be sufficient to accommodate *pir* in as-yet unsampled lineages of the *P(vinckeia)* and *P(plasmodium)* subgenera. To test this, we assembled transcriptomes from RNAseq data generated previously from *Plasmodium*-infected blood taken from two lemur species, *Indri Indri* and *Propithecus diadema* (Larsen *et al*, 2016) (see Materials and Methods, Data files 4 and 5]. Phylogenomic analysis (see Figure S3a) showed that these lemur *Plasmodium* belong to a new lineage that is placed robustly in the phylogeny (ML bootstrap = 99, Bayesian posterior probability = 1), bisecting *P(vinckeia)* and *P(plasmodium)*. We generated Hidden Markov Models (HMM) for each *pir* subfamily (see Data file 6) and used these to categorize all *pirs* among the lemur malaria transcripts. A maximum likelihood phylogeny was estimated with IQTREE2 including the additional lemur malaria *pir* sequences. Lemur malaria *pir* fall within one of the subfamilies characterised here (Figure S3b), usually in agreement with their HMM designation (Figure S3c).

Both *I. indri* and *P. diadema* transcriptomes contain two transcripts most closely related to *pirC1* (i.e. PEXPD_2939316, PEXII_13858233, PEXPD 6773218, PEXII 5438454), further reinforcing the ubiquitous conservation of this locus, and/or its high abundance among bloodstream transcripts. Similarly, several co-orthologs to *pirO1* was observed in I. indri (i.e. PEXII_14251065, PEXII_6556395, PEXII_9023310). Consistent with the position of these lemur malaria parasites at the base *P(plasmodium)*, several of their pir sequences are positioned at the base of pir subfamilies. For example, PEXII_5543448 (*pirM*), PEXII_2114850 (*pirT*, sister to PocGH01_00202100) and PEXII_10236283 (*pirV*, sister to PocGH01_00125800). Transcripts from the parasite infecting *I. indri* suggest that further sampling of malaria parasite species will show that clades currently unique to, or dominated by, *P. ovale*, are not recent *P. ovale* expansions but, in fact, cosmopolitan like other sub-families. For instance, PEXII_2891700, PEXII_7504356 cluster among otherwise *P. ovale*-dominated clades of *pirK*, while PEXII_7818345 clusters within *pirL*, and is sister to PGO_003865, perhaps suggesting a conserved ortholog).

Our analysis of *pir* sequences from an uncharacterised malaria lineage in lemur blood transcriptomes showed that BLASTp could identify *pir* sequences reliably but was not sufficient to precisely place them in a subfamily. HMMER performs better (correctly placing 13/17), but often closely related subfamilies (e.g. *pirJ/K/L*) have approximately equal similarity scores and the result can be ambiguous. These lemur blood transcriptomes are typical in that parasite sequences are often partial and quite short. This, coupled with the probably authentic basal placement of lemur malaria *pir* (e.g *pirT/V*), means that it can be genuinely challenging to unambiguously place a sequence. For example, PEXII_5543448 clusters at the base of *pirM* in the phylogeny but was placed in *pirL* by HMMER, perhaps because but it is not representative of the derived *P. coatneyi/P. knowlesi* sequences that dominate the *pirM* alignment. PEXII_16455210 clusters in *pirP* in the phylogeny but was placed in *pirL/J* (both with a low score), probably because the sequence is too short (68 aa) to be distinctive. We have provided a suite of HMMs describing *pir* subfamilies (Data File 6) for the identification of *pir* in future data sets. However, it is likely that sequencing similarity searches will struggle to correctly place short sequences (<150 aa) and if that fragment comes from the distal-conserved domain, several subfamilies may be equally plausible. Ultimately, while phylogenetic analysis will be required for definitive answers, this application demonstrates that our nomenclature can be used to designate *pir* in any *Plasmodium* genome where they occur, even if not yet characterized.

**Reconciliation analysis is robust to phylogenetic uncertainty in the gene tree**

The numbers of gene duplications and losses inferred by reconciliation analysis are dependent on both gene and species tree topologies. Species tree topology is a well attested consensus (see SI Methods). Gene tree topology, in contrast, contains substantial uncertainty derived from systematic error in the character set, as evidenced by the low bootstrap values throughout Figure 1. Such phylogenetic uncertainty is natural but should be considered when inferring gene family evolution. We examined the effect of systematic error by reconciling 100 bootstrapped gene trees and comparing the distribution of event numbers in these sub-optimal trees with the maximum likelihood consensus shown in Figure 1 (see Figure S4, Data File 7). Figure S4 shows the range in duplication and loss event number for each terminal and ancestral node within the species tree, when reconciled with the bootstrapped gene trees. The overall trends in these ranges is not different to the trends established by the optimal topology (red dots). Thus, the greatest number of gene duplications are inferred at, first, node n18 (i.e. the ancestor of *P(plasmodium)*) and, second, node n12 (i.e. the ancestor of species *P. vivax* to *P. knowlesi* inclusive). The greatest number of loss events are placed at node 6 (i.e. the ancestor of *P. knowlesi, P. coatneyi, P. fragile* and *P. inui*), as well as in *P. malariae* and *P. gonderi*. These results are robust even when variation in sub-optimal gene trees is considered.

**Gene relics in the *P. inui* genome**

Annotated *pir* genes are relatively rare in the *P. inui* genome. There are 10 *pir* sequences that are sufficiently long and intact to include in the phylogeny. As described elsewhere, these include orthologs of well conserved lineages (*pirC1, pirC2, pirO1*; see Table S3). Five sequences belong to *pirI* (C922_01458, C922_03990, C922_05410, C922_04423, C922_05799). Of these ten, three appear to be partial pseudogenes; two are copies of *pirC1* (C922_05050, C922_05571), the other is a copy of C922_05799 (C922_05240). To explore the hypothesis that this meagre *pir* repertoire represents a secondary reduction, the consequence of a genome-wide contraction relative to the ancestral state, (and not a failure to identity *pir* genes), we searched the *P. inui* genome sequence using tBLASTn with all PIR sequences as queries. This revealed a further 8 loci with significant homology to a *pir* but no annotated gene model (see table below). None of these locations contained a full-length and intact gene, so we consider these to be gene relics. Notably, the closest relative of all eight sequences in *P. cynomolgi* is PcyM_1282400, which is sister lineage to C922_03990/C922_05410; this indicates that all gene relics are within-species paralogs of this *pirI* gene. It is also clear that the P. inui genome sequence really does not contain a typical *pir* complement. A similar exercise similarly shows that absence of pir genes in *P. fragile* is real.

| Query (position/bp) | Best match  *P. cynomolgi* M | Sub-family | Align Length | Query Coverage | E-Value | Score | Identity |
| --- | --- | --- | --- | --- | --- | --- | --- |
| 2132018 | PcyM_1282400 | pirI | 284 | 0.969 | 3.18E-22 | 105.0 | 0.690 |
| 5613964 | PcyM_1282400 | pirI | 176 | 0.804 | 4.19E-38 | 156.4 | 0.795 |
| 16302290 | PcyM_1282400 | pirI | 281 | 0.966 | 2.61E-42 | 170.8 | 0.737 |
| 24335288 | PcyM_1282400 | pirI | 249 | 0.428 | 4.17E-18 | 91.5 | 0.687 |
| 24979937 | PcyM_1282400 | pirI | 241 | 0.430 | 2.23E-21 | 102.3 | 0.693 |
| 25330886 | PcyM_1282400 | pirI | 265 | 0.530 | 3.12E-25 | 115.8 | 0.702 |
| 26723060 | PcyM_1282400 | pirI | 213 | 0.411 | 1.29E-17 | 89.7 | 0.700 |
| 26724033 | PcyM_1282400 | pirI | 250 | 0.943 | 5.16E-19 | 93.3 | 0.688 |

**Interspecies differences in *pir* diversity do not depend on genome version**

The low numbers of *pir* genes in *P. inui* and *P. fragile* genomes sequences may reflect the effort that has been put into their sequencing. Both genome sequences were generated using short-read technology in 2014 and 2016, respectively, and have not been improved with more recent technologies (Genbank references: GCA_000524495.1 for *P. inui* and GCA_000956335.1 for *P. fragile*). To test this, we counted the number of pir genes in early genome versions for species with well-developed genome sequences using BLASTn. In the current PlasmoDB release (68), *P. knowlesi* has 68 genes in 3 subfamilies; in release 10, it had 177 genes (in 3 subfamilies). *P. vivax* has 287 genes in 18 subfamilies; it had 289 genes (in 18 subfamilies) in database release 5 and 336 genes (in 18 subfamilies) in release 10. *P. ovale* has 1269 genes in 22 subfamilies; it had 1407 genes (in 22 subfamilies) in database release 30. Finally, *P. yoelli* has 766 genes in 2 subfamilies; it had 905 genes (in 2 subfamilies) in database release 5 and 794 genes (in 2 subfamilies) in release 10. Thus, the number of annotated *pir* genes in a genome sequence does not increase as *Plasmodium* genome sequences have improved (in fact, it generally falls as partial gene models are combined). Neither does the number of recognised subfamilies change. For these species, their early draft genome versions capture the diversity of *pir* genes that we see in more mature versions. Therefore, we can be confident that the dearth of *pir* genes in the *P. fragile* and *P. inui* genome sequences is not due to the relatively early stage of their sequencing.
